# T-cell microvilli simulations show operation near packing limit and impact on antigen recognition

**DOI:** 10.1101/2021.11.24.469916

**Authors:** Jonathan Morgan, Johannes Pettmann, Omer Dushek, Alan E. Lindsay

**Affiliations:** Dept. of Applied and Computational Mathematics and Statistics, University of Notre Dame, Notre Dame, IN, 46556, USA; Sir William Dunn School of Pathology, University of Oxford, Oxford, OX1 3RE, UK; Biophysics Graduate Program, University of Notre Dame, IN, 46556, USA

**Keywords:** Immune signaling, T-cell activation, Microvilli dynamics

## Abstract

T-cells are immune cells that continuously scan for foreign-derived antigens on the surfaces of nearly all cells, termed antigen presenting cells (APCs). They do this by dynamically extending numerous protrusions called microvilli (MV) that contain T-cell receptors (TCRs) towards the APC surface in order to scan for antigens. The number, size, and dynamics of these MV, and the complex multi-scale topography that results, play a yet unknown role in antigen recognition. We develop an anatomically informed model of the T-cell/APC interface to elucidate the role of MV dynamics in antigen sensitivity and discrimination. We find that MV surveillance reduces antigen sensitivity compared to a completely flat interface unless MV are stabilized in an antigen-dependent manner and find that MV have only a modest impact on antigen discrimination. The model highlights that MV contacts optimise the competing demands of fast scanning speeds of the APC surface with antigen sensitivity and that T-cells operate their MV near the interface packing limit. Finally, we find that observed MV contact lifetimes can be largely influenced by conditions in the T-cell/APC interface with these lifetimes often being longer than the simulation or experimental observation period. The work highlights the role of MV in antigen recognition.

**Significance Statement:** T-cells search for foreign-derived antigens on the surface of antigen presenting cells (APC) by dynamically extending tubular protrusions called microvilli (MV) which form membrane close-contacts. Although known for decades, their role in antigen recognition remains unclear. Guided by recent experiments, we built an anatomically informed stochastic model of MV scanning and compared with a topologically flat interface. We find that MV scanning modestly impacts antigen discrimination, yet it enables T-cells to balance the competing effects of maintaining sensitivity while conducting rapid APC surveillance. The model can reconcile discrepancies in observed MV lifetimes and demonstrates that observed area coverage fractions correspond to geometric packing limits. Our work suggests that MVs promote positive signaling outcomes despite anatomical constraints to close contact formation.

---

T-cells are immune cells that continuously search for the molecular signatures (antigens) of pathogens and upon antigen recognition, can initiate adaptive immune response. T-cells recognize antigen through binding of their T-cell receptor (TCR) to antigens presented by antigen presenting cells (APCs). Interestingly, T-cells search for antigen on APCs via small tubular membrane protrusions, termed microvilli (MVs). The existence of MV has been known for decades, yet the function of these structures is largely unknown.

Microvilli are generally described to be ≈50–400nm wide and micrometer long structures (1). They first became apparent when imaging T-cells by electron microscopy (EM) (2–4). Due to their small size, conventional light microscopy cannot resolve them. However, recent developments have allowed imaging of microvilli on cells (5, 6), greatly increasing our understanding of their dynamics in homeostasis or during antigen recognition. While they only cover about 40% of the interface at a given time, through their dynamic movements they are able to scan 98% of the interface with an APC within physiological dwell times of T-cell/APC contacts *in vivo* (5, 7–9). The lifetime of an individual MVs is short when it does not recognise antigen (≈8 seconds) but can increase upon antigen recognition with reported lifetimes ranging from 9 seconds to 18 minutes (5, 6). Super-resolution microscopy has revealed that MV tips are enriched in TCR and adhesion molecules (4, 10), consistent with their role in forming the small close contacts required for TCR/antigen binding (11). One hypothesis is that MVs are a necessity of the biophysical constraints of the T-cell/APC interface (1). Both the surface of APCs and T-cells are covered with glycoproteins, such as the abundant CD45 on T-cells, generating a thick glycocalyx that causes steric and electrostatic repulsion (12–14). As both TCRs and antigen are much shorter in comparison, the penetration of this matrix is required for the formation of close membrane contacts and binding to occur (6, 14).

The potential role of MVs in antigen recognition is unclear. The process of antigen recognition is marked by rapid recognition speeds with single-molecule sensitivity to agonist ligands. For example, CD4^+^ T-cells can exhibit Ca^2+^ signaling, a precursor of T-cell activation, when stimulated with a single molecule of foreign antigen (15). Similarly, CD8^+^ T-cells have been shown to recognize as few as 3 molecules of a foreign antigen (16). This single-digit molecule sensitivity is particularly remarkable given how short lived TCR/antigen interactions are. TCRs bind foreign antigen with typical bond lifetimes in the range of 1–10 seconds (17). Moreover, this level of sensitivity is achieved with a high speed, since T-cells have been shown to exhibit Ca^2+^ signaling in as little as 7 s upon antigen binding (18). Intuitively, MVs would be expected to decrease the sensitivity when compared to a flat interface as they only contact 40% of the surface at a given time.

Lastly, T-cells are able to amplify small differences in antigen affinity into large differences in their responses (19). Furthermore, T-cells can ignore APCs that express a high abundance of self-derived antigen while retaining sensitivity to only a few foreign antigen, even in cases where they may be chemically similar (20). We have previously speculated that small contacts, such as the ones formed by MVs, could improve the antigen discrimination (21). In this theory, creating small domains improves signal/noise, effectively implementing kinetic proofreading on a topological level. Furthermore, antigen discrimination was shown to be improved by a biophysical phenomenon that can increase the lifetime of foreign antigens (catch bond), while decreasing the lifetime of self antigens (slip bond) (22–25). In this model, MV may act as the source of the bond stress (1).

In the present work, we develop a new computational model that combines stochastic topography induced by MV surveillance of the APC surface together with TCR/antigen interactions. We investigate the effect of MVs on sensitivity, speed, and antigen discrimination when compared to a hypothetical flat interface. We find that the sensitivity and speed are decreased with MVs, while the discrimination is largely unchanged. However, when introducing signalling-dependent MV stabilization, sensitivity and speed are markedly improved. Lastly, we show that MV scanning operates near the packing limit of MVs in the contact interface and that discrepancies in the experimentally measured MV lifetimes are likely a result of censored data collection. Together, these results support the hypothesis that MVs have evolved out of biophysical constraints, rather than to improve antigen recognition or discrimination.

## 1. Results

### Model

In our model of T-cell activation, TCRs and antigen interact only via close contacts formed by MV tips which form (with rate *k*_a_) into available open T-cell/APC space and are subject to removal (with rate *k*_d_) (Figure 1: See section 3 for details). Rate parameters are determined using experimental observations of surface area coverage by MV tips and packing simulations to find an approximate maximum number of contacts that can be placed in a circular, T-cell/APC interface of diameter of 5*µ*m (5). TCR/antigen interactions are described by a kinetic proofreading process (26) with a fixed number of steps (*N*), and transition rate (*k*_t_), obtained from recent experimental work (19). Signaling TCR/antigen pairs are integrated equally over all spatial regions with T-cell activation defined by the accumulated signal surpassing a pre-set threshold. MV contacts housing an activated TCR may become *stabilized* (with rate *k*_s_) extending their lifetime (Figure 1) with removal only possible when there are no activated TCRs within the contact. To assess discrimination across model and parameter regimes, we utilize a *discrimination power* (*γ*) to quantify the amplification of T-cell response from variations in dissociation rates (*k*_off_). The factor *γ* is determined as the slope of a linear fit of *P*_15_ (antigen population necessary to elicit a 15% chance of T-cell activation) with *k*_off_ (19). Larger values of *γ* imply better discrimination (Figure 2A-B and supplement).

**Fig. 1.**
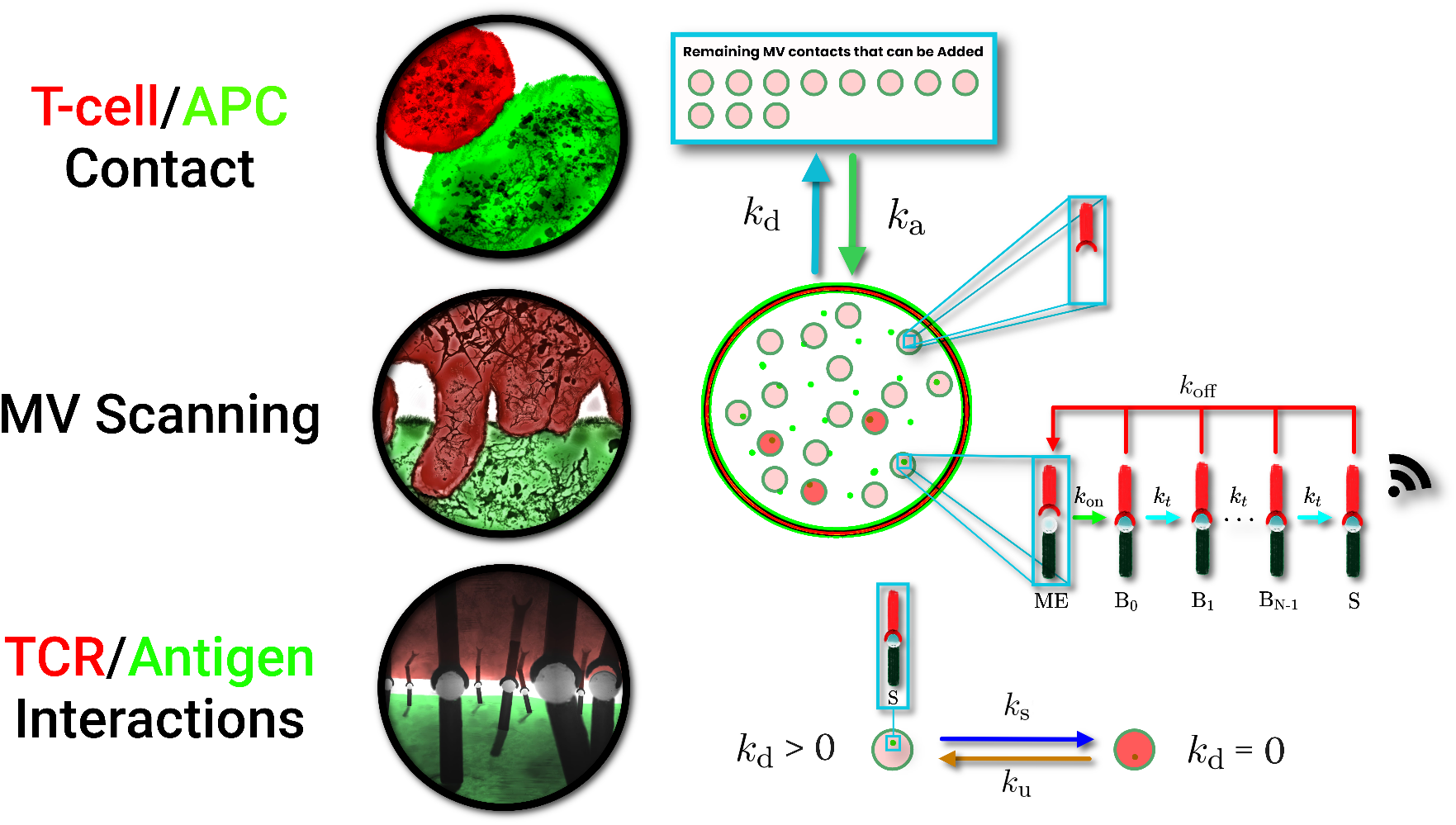
Model overview. (Left) Illustrations of the three layers involved in the stochastic MV scanning model. TCR/antigen interactions are facilitated by close contacts formed on the tips of MV. All interactions take place in the T-cell/APC interface. (Right) Illustrative description of the simulated dynamics of the T-cell/APC interface. Close contacts by MV are formed at rate *k*_a_ and are removed with rate *k*_d_, given that the contact has not been stabilized. TCRs and antigen interact within the close contact zones according to the displayed kinetic proofreading mechanism. MV contacts that contain an activated TCR within the contact boundary stabilize with rate *k*_s_. Stabilized MV contacts can not be removed, i.e., *k*_d_ = 0. Instead, these contacts can only be destabilized with rate *k*_u_, given that there are no activated TCRs within the MV contact boundary.

**Fig. 2.**
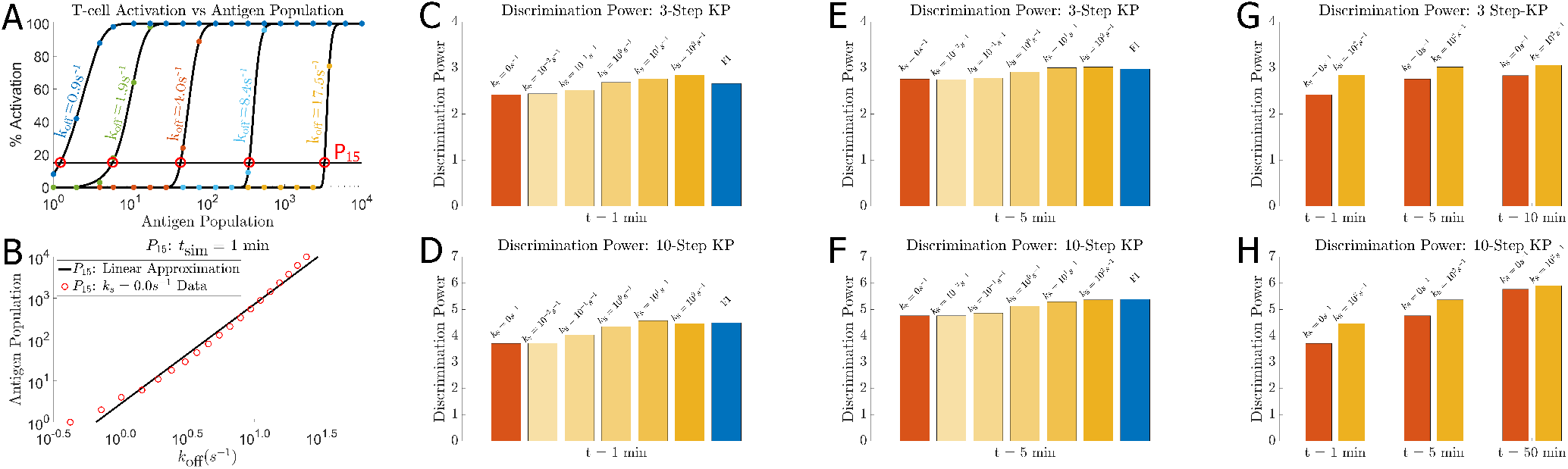
MV scanning and discrimination power. **A** Proportion of activated T-cells across antigen numbers. The quantity *P*_15_ is the antigen number required to produce at least a 15% chance of activation. **B** Determination of the discrimination power *γ*. **C–H** Comparisons of the measured discrimination powers among several models. Results shown are of models with either N=3 or N=10 kinetic proofreading steps. Times shown at the bottom of each graph indicate the simulated duration of the T-cell/APC contact.**C–D** Discrimination powers for 3-step (**C**) and a 10-step (**D**) kinetic proofreading steps, simulated for 1 minute of T-cell/APC contact time. **E–F** Discrimination powers for 3-step (**C**) and a 10-step (**D**) kinetic proofreading steps, simulated for 5 minutes of T-cell/APC contact time. **F. G–H** Discrimination powers between MV models for different T-cell/APC interface times with 3-step (**G**) and 10-step kinetic proofreading (**H**).

### MV scanning modestly affects antigen discrimination

We measured discrimination power (*γ*) over three models (flat topology, MV scanning with both strong stabilization and no stabilization) and across parameter regimes (*N* = 3, 10 proofreading steps and simulated T-cell/APC contact times *t*_sim_ = 1, 5, 10 mins). Across these scenarios, we found that *γ* did not vary over biologically significant ranges, suggesting that MV scanning neither significantly hampers nor greatly improves antigen discrimination (Figure 2). This is despite MV scanning only facilitating instantaneous contact between 40% of the opposing surfaces, therefore generally reducing the probability of T-cell activation at a given TCR/antigen dissociation rate when compared to a flat topology.

The most significant modulation in discrimination power was observed in the case of short lived T-cell/APC contacts (*t*_sim_ = 1 min) with strong MV stabilization (*k*_s_ = 100*s*^−1^) (Figure 2G–H). Clearly stabilization can partially negate some constraints imposed by MV scanning by extending MV lifetimes, thus allowing sufficient time for productive TCR/antigen interactions to occur. To explore the relevance of short contact times, and to better understand model predictions of antigen discrimination, we derived potency curves (*P*_15_) from a mean field approximation of the stochastic model. We found that in small populations of antigen, or for short T-cell/APC contact times (*t*_sim_), stochastic effects can play an outsize role such that stabilization of MV contacts results in significant improvement to sensitivity (see supplement).

### MV stabilization optimizes scanning speed and agonist sensitivity

In addition to exploring the discrimination powers of MV and flat interface models, we further sought to investigate the role of MV contact stabilization on T-cell activation time and antigen sensitivity. In the structure of our model, larger values of *k*_d_ correspond to faster surveillance of the APC surface (Model and Methods) which results in a more rapid cumulative survey of the interface (Figure 3E). However, rapid scanning by MV further constrains antigen recognition by increasing the probability that a contact is removed before a TCR can become activated. To explore this trade off, we simulate the T-cell/APC interface with a single antigen and display in Figure 3A–D the average T-cell activation times and the percentage of activated T-cells in a population over a varying *k*_d_ and *k*_off_. In the absence of MV contact stabilization, we observe that T-cell activation times are approximately homogeneous in regions where T-cell activation is likely (Figure 3A). In such regions, the probability of T-cell activation is also similar (Figure 3B). For example, changing *k*_d_ from *k*_d_ = 10^−3^ to *k*_d_ = 10^−1^ yields little variation in T-cell activation times and probabilities, given a strong agonist (*k*_off_ = 10^−2^). Without MV stabilization, increasing *k*_d_ increases MV scanning speeds, but also decreases the probability of T-cell activation. The same is not always true for models with MV contact stabilization. For a model with contact stabilization, there are cases where increasing *k*_d_ increases both, MV scanning speed and the probability of T-cell activation (Figure 3C and D). This is due to MV stabilization reducing the probability of contact removal following TCR activation, i.e., reducing the probability of having to search for an agonist again, once an agonist is found. Additionally, the fastest T-cell activation times and the highest probability of activation occur in this region of higher scanning speeds. This region of optimal *k*_d_ is nearly completely absent in models without contact stabilization.

**Fig. 3.**
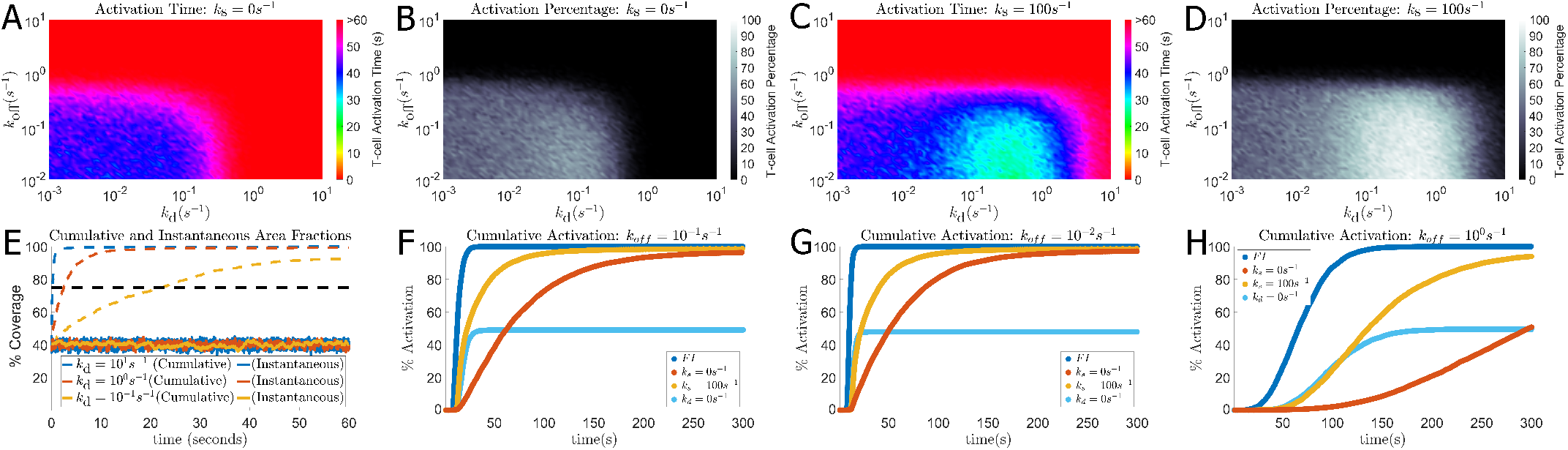
Results for *N* = 3 kinetic proofreading steps and a T-cell/APC interface time of *t*_sim_ = 60*s*. (**A**) Average T-cell activation time with a single agonist in the interface over a varying *k*_d_ and *k*_off_ without stabilization (*k*_s_ = 0*s*^−1^). Simulations that did not activate were weighted in the average by the total simulation time, *t*_sim_ = 60*s*. (**B**) Percentage of T-cells that are activated under the same conditions as (**A**). (**C**-**D**): Similar to (**A**-**B**) with strong stabilization *k*_s_ = 100*s*^−1^. (**E**) Cumulative (dashed) and instantaneous (solid) scanning of an opposing surface in simulations with MV scanning and no stabilization, over a varying *k*_d_ and varying *k*_a_ (Eq. (3)). (**F–H**) Cumulative activation percentage in a population of T-cells given an increasing TCR/antigen dissociation rate. Each simulation contained a single agonist in the T-cell/APC interface and had an allowed interface time of *t*_sim_ = 300*s*.

In Figure 3A–D, the instantaneous area fraction is maintained at 40% for all *k*_d_, which is accomplished by setting *k*_a_ = (14/3)*k*_d_ (Model and Methods). Such a fraction has been observed experimentally when imaging microvilli (5). In Figure 3E we show how this relationship is maintained in individual simulations. The instantaneous scanning fraction remains approximately within the bounds of 40 ± 5% while the cumulative scanning speed can increase or decrease drastically given an increase or decrease in *k*_d_, respectively. The extreme case *k*_d_ → 0*s*^−1^, which implies *k*_a_ → 0*s*^−1^ by our definition of *k*_a_, reflects a variant of the flat interface model where 40% of the T-cell surface is covered in TCR clusters (leftmost regions of Figure 3A–D).

We further explore the intricacies of T-cell activation in our model by observing the cumulative T-cell activation in a population over a period of time, *t*_sim_. We simulate four separate models corresponding to a flat interface, MV scanning, MV scanning and contact stabilization, and an additional model in which *k*_d_ = 0*s*^−1^ and *k*_a_ is constant. The additional model here is representative of a flat interface in which only certain regions are densely packed with TCRs, i.e., a model with TCR clusters. In Figure 3F–G we show the results of such simulations for models with *N* = 3 kinetic proofreading steps simulated for *t*_sim_ = 300*s* and a single agonist in the interface with a TCR/antigen dissociation rate of *k*_off_ = 10^−2^*s*^−1^ (Figure 3F), *k*_off_ = 10^−1^*s*^−1^ (Figure 3G), and *k*_off_ = 10^0^*s*^−1^ (Figure 3H). As would be expected, the flat interface model (dark blue), where the entire surface is assumed to be densely packed with TCRs, achieves the highest percentage of activation in the shortest amount of time. This is due to the absence of a search process for agonists, as well as every agonist being accessible to TCRs in the T-cell/APC interface. Additionally, we observe that the flat interface model with localized TCR clusters (light blue) is just a scaled cumulative activation profile of the densely packed flat interface model. Hence, we can expect that any flat interface model with differing distributions of TCR clusters will be a scaled variant of the densely packed flat interface model, in which the magnitude of the scaling is approximately the fraction of surface area covered by said clusters. However, it is important to note that this is a model dependent phenomenon and does not take into account the role of antigen diffusion. In the MV scanning models, we observe faster T-cell activation in a population when MV contacts can be stabilized (gold) versus MV scanning without contact stabilization (red). This is unsurprising given the results shown in Figure 3A–D, as we found that T-cells with stabilized MV contacts, opposed to those without, activated the fastest when there was only a single agonist in the interface. Since faster scanning speeds serve little to no purpose in large antigen populations, this optimal scanning speed is present in only smaller populations of antigen.

### Model explains observed discrepancies in MV contact lifetimes

Observed microvilli lifetimes (*τ*) in experiments have varied significantly. For example, Cai et al. observed MV lifetimes of *τ* ≈ 7.7*s* in the absence of agonists and *τ* ≈ 8.9*s* in the presence of agonists (5). In contrast, another group studying membrane protrusions on T-cells observed much longer MV lifetimes, on the range *τ* ≈ 60*s* when agonists were not present and *τ* ≈ 18 minutes when agonists were present (6). To understand the plausibility of such a discrepancy, we use our model with MV stabilization to explore lifetimes over a range of varying dissociation rates and antigen populations. We find that MV lifetimes can easily surpass our simulation time when large numbers of agonists with low TCR/antigen off-rates are used (Figure 4A). This is expected, given that the probability of finding a TCR/antigen complex in a state where the stabilized MV is capable of being destabilized is *t*_kp_/(*t*_kp_ + *t*_S_), and is the steady state probability that the TCR is not in an activated state. Here, *t*_kp_ is the average time it takes a TCR/antigen pair to bind and undergo all *N* kinetic proofreading modifications (supplement), including all dissociation events, before becoming activated, and *t*_S_ = 1*/k*_off_ is the average time a TCR stays activated once it reaches the final, signaling-competent stage. As *k*_off_ decreases, *t*_kp_ decreases and *t*_S_ increases for a given antigen capable of binding TCRs. Hence, the probability of finding the antigen not bound to an activated TCR decreases. When there are *n* antigen covered by an individual stabilized MV contact, this probability becomes (*t*_kp_/(*t*_kp_ + *t*_*S*_))^*n*^ and tends towards zero as *n* grows large. The small probability of finding all antigen not bound to an activated TCR also indicates that the MV contacts spends very little time in a state capable of being destabilized (Model and Methods). Conversely, if there are very few antigen, or the antigen are weak agonists, MV stabilization may be a rare event. In these conditions, the average MV lifetime *τ* would be approximately *τ* = 1*/k*_d_, which is 7*s* in our parameterization. This suggests that MV lifetimes are not constant and are determined by the conditions in the T-cell/APC interface.

**Fig. 4.**
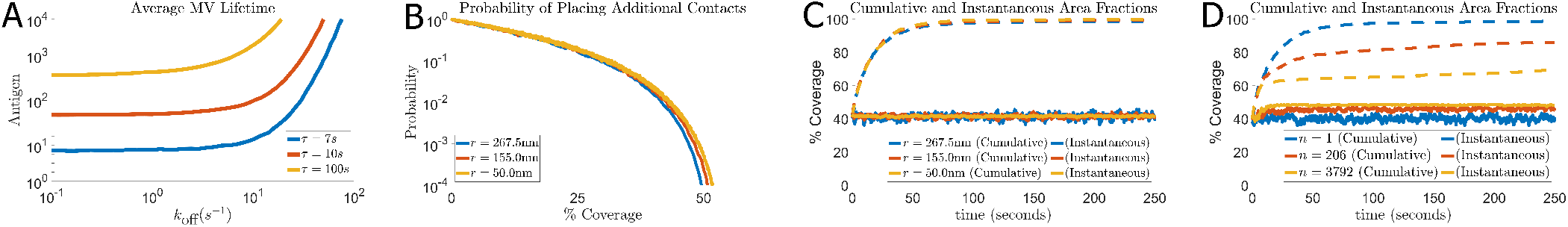
Stochastic simulations of MV placement and dynamics in model with stabilization. (**A**): Contour plot of average MV lifetimes (*τ*) over antigen population (*n*) and TCR/antigen disassociation rates (*k*_off_). (**B**): Likelihood of successful MV tip placement into open space of the T-cell/APC interface against MV coverage fraction. (**C**): Instantaneous (solid line) and cumulative (dashed line) scanning area fractions in the absence of agonist antigen for several MV tip radii. (**D**): Similar to panel (**C**) but curves represent different numbers (*n*) of agonist antigen present.

### Observations of MV dynamics indicate operation near theoretical packing limits

We explored the dynamics of other MV radii (50nm, 155nm, 268nm) in our model in which span the range of widths reported from light and electron microscopy measurements (3, 4, 6). When we ran MV packing simulations, where MV cannot be removed and locations for additional MV are randomly chosen, but overlap between MV is forbidden, we found that the maximum packing fraction converged to approximately 50% coverage (Figure 4B). While the exact packing fraction may depend on the MV radius, the practical packing fraction (i.e. what can be achieved by random placement and without replacing MV) did not vary strongly. This indicates that the 40 ± 5% instantaneous area coverage observed when imaging T-cells interacting with antigen presenting cells (5) may be a result of a practical packing limit of MV. Furthermore, we show that our stochastic model of MV scanning can achieve similar cumulative scanning fractions to that of observations by Cai et al.(5), as well as maintain the instantaneous area fraction at approximately 40 ± 5% (Figure 4C). We note that varying the MV radius had little effect on the cumulative scanning fractions, with only a slight increase in scanning speed resulting from a smaller MV radii. In addition, previous studies reported a decreased scanning speed of MV when foreign antigen were present on an APC surface (5, 6). Consistent with this, our MV stabilization model can reproduce this observation, where foreign-antigen slow cumulative scanning and increasing numbers of agonists slows scanning speed further (Figure 4D). When there are very large populations of strong agonists in the T-cell/APC interface, the instantaneous scanning fraction approaches the maximum packing fraction and the cumulative scanning of MV halts. This is the result of MV contacts being fixed in place by stabilization for the entirety of a simulation.

## 2. Discussion

Microvilli on T-cells have been described for decades, yet their function remains enigmatic. We developed a new stochastic model of T-cell/APC engagement which reveals that MV scanning and the stabilization of MV contacts works jointly to maximize the probability and speed of T-cell activation by optimizing both sensitivity to agonists and MV scanning speeds of the opposing APC surface. In particular, MV scanning with stabilization can significantly improve the sensitivity of T-cells to small populations of agonists.

We sought to identify how antigen discrimination might be modulated given recent observations that only 40% of the T-cell/APC interface are in physical contact. Over a wide range of modeling scenarios and parameters regimes, we found that antigen discrimination was neither diminished nor significantly improved by the action of MV. It is therefore possible that microvilli-based scanning mode has not evolved to improve discrimination, but rather due to the biophysical constrains preventing formation of large flat interfaces. Specifically, the glycocalyces of T-cells and APCs are expected to prevent large contact zones due to steric, electrostatic, and entropic effects (1). In addition, hydrodynamic effects have been suggested as limits on the speed of contact formation - an effect that becomes more significant with increasing contact size (27). Our model yields a wide range of MV lifetimes (5, 6) and suggests that MV contact lifetimes may be highly dynamic in nature. The average MV contact lifetime can be highly dependent on the conditions in the T-cell/APC interface. It is not unusual to have MV lifetimes that can far exceed the T-cell/APC interface time when there are medium to large populations of agonists present. When MV contact lifetimes exceed the T-cell/APC interface time, or recording period, the average of these MV lifetimes also becomes dependent on the length of time over which the data is collected. Finally, we point out that the 40 ± 5% instantaneous area coverage by MV may be close to a packing limit of MV contacts in the T-cell/APC interface. Our results show that as more contacts become stabilized by TCR interactions with agonists, MV scanning slows significantly and the instantaneous area coverage will approach this limit. Further research may reveal if they have primarily evolved to improve discrimination or due to the biophysical constrains preventing a flat interface.

## 3. Model and Methods

The basis of our stochastic model is the kinetic proofreading (KP) mechanism (26) that proposes a productive immune signal can arise from TCR/antigen interactions only after a sequence of intermediate binding steps have occurred. Each intermediate step is irreversible, such that upon TCR/antigen dissociation the TCR reverts to its original unphosphorylated state before a new antigen binds. KP predicts that T-cell activation is linked to the lifetime (1/*k*_off_) of the TCR/antigen complex (26). An outcome of this mechanism is that a time delay is induced between an initial binding event and eventual signaling, such that only long-lived TCR/antigen complexes progress into a signaling state. With multiple steps, kinetic proofreading can discriminate with high accuracy between antigens of different TCR/antigen dissociation rates (*k*_off_).

Our model consists of *N* intermediary states {*B*_1_, …, *B*_*N*_} with transition rate *k*_t_. The number of kinetic proofreading steps was set to *N* = 3 (unless stated otherwise) and the transition rate set to *k*_t_ = 1.0*s*^−1^, due to recent experimental work (19). The rate *k*_on_ is assumed to be the product of the bimolecular on-rate and the 2D radial density of TCRs (*k*_on_ = 100*s*^−1^), and is taken to be relatively fast compared to other model parameters. The TCR concentration is assumed to be relatively high, i.e., the flat interface and MV tips are densely packed with TCRs. If the model relies on multiple rebinding steps (TCR+Antigen → *B*_1_) before a TCR can become activated, then on-rates can greatly influence antigen sensitivity, as observed experimentally (28–30). In such a case, antigen with fast on-rates led to T-cell activation while those with slower binding rates did not. An assumption in our model is that antigen discrimination is primarily influenced by the dissociation rate *k*_off_ to the TCR, therefore we chose *k*_on_ to be fast and constant for all antigen. Other evidence for fast TCR/antigen binding was seen in Cai et al. (5) who observed that TCR recognition occurred at the same time as surface deformations (MV) contacted the APC. Other studies have suggested that describing TCR/antigen binding with an on-rate may be inappropriate (31) with their results indicating that binding may occur after a minimal encounter duration (*t*_on_ < 1ms).

In each of our T-cell activation models, we assume the T-cell/APC interface, or immune synapse (IS), to be circular with radius *r*_IS_ ≈ 5*µm* (5).

### Flat Interface Model

Our idealized flat topology model is the stochastic implementation of the kinetic proofreading model proposed by McKeithan (26). There are no structures such as MV, and all antigen within the T-cell/APC interface are accessible to TCRs. The assumption that the T-cell surface is densely packed with TCRs implies that when the contact is formed, every antigen present in the contact area is close enough to bind to one or more TCRs. Utilizing this assumption, we consider only the discrete population of antigen in binding interactions, rather than concentrations of both species (Fig. 1A and B). This idealized model serves as an upper bound of antigen accessibility to TCRs and is utilized due to its likeness to the deterministic mass action model.

The rate at which signal accumulates from individual activated TCRs, as well as the threshold of total signal required for T-cell activation, is chosen arbitrarily such that antigen discrimination is possible within experimentally observed dissociation times. As reported by Iezzi et al. and Huppa et al., TCR/antigen complexes can experience lifetimes as short as 0.1*s* to 10*s* (32–34). The model includes an accumulated signal *X*(*t*) to the T-cell with dynamics given by

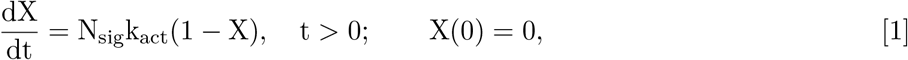

where *k*_act_ is the signaling rate and *N*_sig_ is the number of TCR/antigen pairs in a signaling state. To obtain a digital like response, we define a T-cell to be activated when *X > X*_act_, where *X*_act_ is the global signal required for activation, and the associated time of activation is min_*t*>0_ {*X*(*t*) = *X*_act_}. The dynamics of this function resemble experimental observations of signal accumulation in T-cells (35, 36). The parameter values *k*_act_ = 0.2*s*^−1^ and *X*_act_ = 0.90 were chosen such that T-cell activation is unlikely to occur with a single visit to a signaling state by a weak agonist while also controlling the range of dissociation rates where discrimination is likely to occur. Signal accumulation occurs globally, meaning that regardless of the spatial location of activated TCR/antigen pairs in the IS, they contribute equally to the total accumulated signal. Similar behavior has been observed in experiments utilizing photoactivation of micro-scale regions of the IS, where it was found that localized signals propagate away from stimulated regions indicating that signals from upstream events act cooperatively to elicit a generalized calcium response (18).

### MV Scanning and Stabilization

We extend the kinetic proofreading model to incorporate MV dynamics, with the assumption that TCR/antigen interactions only occur when the MV provides a close contact. MV contacts are represented by non-overlapping circular regions of fixed radii, *r*_MV_, and are placed randomly in the IS with rate *k*_a_. When a contact is added to the interface, its position is fixed until the MV is removed. If the MV contact encompasses one or more antigen, then TCR/antigen interactions can occur according to Figs. 1C and 1D. Each MV contact is capable of having multiple simultaneously activated TCRs, up to the number of antigen covered. If the MV contact does not cover any antigen, then the contact can only be removed. In the MV scanning model without contact stabilization contacts are removed with rate *k*_d_, regardless of the state of any resident TCRs. This is not the case when *k*_s_ *>* 0, where MV lifetimes can be significantly extended when a TCR becomes activated.

The instantaneous fraction of T-cell surface area covered by MV was reported to be *f* ≈ 40 ± 5% (5). This area fraction range was observed in both the IS and isolated regions of the T-cell, with and without agnostic antigen. The individual MV radius, *r*_MV_ ≈ 267.5*nm*, and the average number of MV contacts, 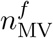 in the T-cell/APC interface are related by

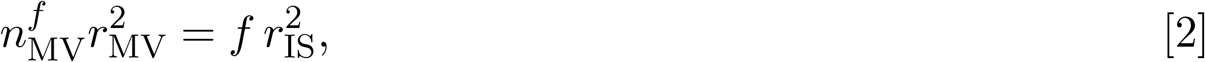

which yields 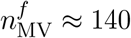 (5). The rates of MV contact placement *k*_a_ and MV contact removal *k*_d_ are related at equilibrium by

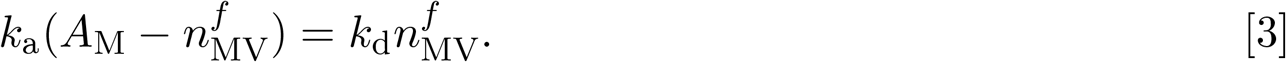

The term *A*_M_ represents the packing limit or maximum number of non-overlapping circular contacts with fixed radius *r*_MV_ that can simultaneously occupy the IS.

In the presence of foreign antigen, extended lifetimes of MV contacts have been observed (5, 37). We reproduce this behavior by extending the dynamics to include a stabilized MV state (asterisk marked states in Fig. 1) in which a stabilized contact must first become destabilized (rate *k*_u_ = 0.03*s*^−1^) before said contact can be removed. The average lifetime of stabilized MV contacts is chosen to exceed the 11.1*s* observed by Cai et al. (5) due to their finding that 25% of stabilized contacts were stable for the entire observation period. We surmise that stabilization is a rapid process induced when a TCR enters an activated state and reflect this in the parameter selection *k*_s_ = 100*s*^−1^. This effectively means that MV become stabilized when antigen binds to the TCR for sufficient duration. However, we also evaluate cases where 0*s*^−1^ < *k*_s_ ≤ 100*s*^−1^ in order to show the influences of an increasing rate of MV stabilization (Fig. 2A–F).

### MV Contact Packing Limits in the IS

To prevent steric clashes between MV in the simulations, we pre-determine possible packing limits using random contact placement. As mentioned previously, *k*_a_ is chosen such that approximately 40 ± 10% of the area is covered by MV contacts at any time and this rate varies with *r*_MV_. For a fixed MV tip radius *r*_MV_, the packing limit is determined by repeated attempts (*T*) at randomly placing non-overlapping contacts in the T-cell/APC interface. The simulation is terminated when the number of placement attempts exceeds a threshold (*T > T*_Max_), after which the final number of MV contacts is recorded. This simulation process is repeated 100 times and the results are averaged to obtain *A*_M_. In Fig. 4B, the results of packing simulations for different MV radii (*r*_MV_) are shown. The vertical axis (1/*T*) approximates the probability of placing an additional MV contact into an open space in the T-cell/APC interface as the number of contacts (*n*_MV_) increases.

**Table 1.** Parameter values used in Figs. 7,8, and 9.

| Parameter | (Unit) | Flat Interface | MV Scanning | MV Stabilization |
| --- | --- | --- | --- | --- |
| $k_a$ | $(s^{-1})$ | N/A | $\frac{2}{3}$ | $\frac{2}{3}$ |
| $k_{on}$ | $(s^{-1})$ | 100 | 100 | 100 |
| $k_t$ | $(s^{-1})$ | 1.0 | 1.0 | 1.0 |
| $k_d$ | $(s^{-1})$ | N/A | $\frac{1}{7}$ | $\frac{1}{7}$ |
| $k_s$ | $(s^{-1})$ | N/A | N/A | $[0, 100]$ |
| $k_u$ | $(s^{-1})$ | N/A | N/A | 0.03 |
| $k_{off}$ | $(s^{-1})$ | $[10^{-1}, 10^{1.5}]$ | $[10^{-1}, 10^{1.5}]$ | $[0.01, 10]$ |
| $k_{act}$ | $(s^{-1})$ | 0.2 | 0.2 | 0.2 |
| $N$ | None | 3 | 3 | 3 |
| $r_{MV}$ | (nm) | N/A | 267.5 | 267.5 |
| $r_{IS}$ | ($\mu m$) | 5.0 | 5.0 | 5.0 |

### Parameters

- *k*_a_: Derived to facilitate a steady state MV contact population such that 40% of the T-cell/APC interface is covered (5).
- *k*_on_: Chosen to be fast to reduce the effects of on-rates in antigen discrimination, observed in Govern et al. (29). Binding was observed on the order of ms in Limozin et al (31).
- *k*_t_: Set according to experimental work by Pettmann et al. (19).
- *k*_d_: The average lifetimes for MV contacts was recorded to be ∼ 20*s* (5).
- *k*_s_: No direct experimental value is known. In Fig:2 we vary *k*_s_ over several values ([0*s*^−1^, 100*s*^−1^]) and find that there is limiting behavior as *k*_s_ ≫ *k*_d_. The most significant effects of contact stabilization are seen when *k*_s_ approaches this limit (2G). Hence, we utilize a maximum stabilization rate of 100*s*^−1^ = *k*_s_ ≫ *k*_d_ = 1/7*s*^−1^.
- *k*_u_: No direct experimental value is known. Set low to replicate experimental observations of long MV lifetimes in the presence of agonists (5).
- *k*_off_: Set such that the values for which we varied *k*_off_ would be similar to dissociation rates reported in the literature (32–34).
- *N* : The number of kinetic proofreading steps used in our model. Set according to experimental work by Pettman et al. (19)
- *r*_MV_: Chosen such that MV widths are relevant to experimental measurements (3, 4, 6). At *r*_MV_ = 267.5*nm*, 140 MV contacts make up approximately 40% coverage in the interface.
- *k*_act_: Chosen arbitrarily with the condition that discrimination occurs within the varied interval of *k*_off_.

## ACKNOWLEDGMENTS

AEL acknowledges support from the National Science Foundation through awards DMS-1815216, DMS-1931705 and DMS-2034787. OD acknowledges the support of a Wellcome Trust Senior Fellowship in Basic Biomedical Sciences (207537/Z/17/Z). JP acknowledges the support of a Wellcome Trust PhD Studentship in Science (203737/Z/16/Z)

