## Supplemental Material for "T-cell microvilli simulations show operation near packing limit and impact on antigen recognition"

Compiled on November 29, 2021

### 1. Mass action kinetic proofreading model

Our formulation of a mass action model follows from the theoretical work proposed by Mckeithan (1). Mckeithan proposed a continuous model to approximate the fraction of activated TCR/pMHC complexes. The model was introduced to better understand the specificity observed in antigen discrimination at the level of a single TCR.

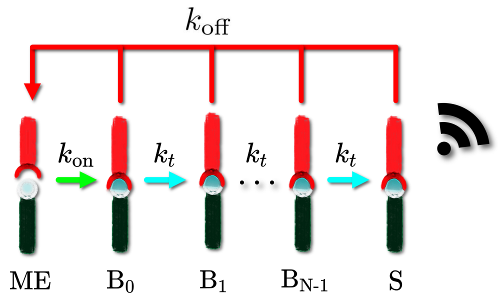


**Fig. 1.** Kinetic Proofreading Model

The fractional steady states of the model depicted in Figure 1 can be computed for arbitrary parameters.

$$\begin{aligned}
\text{ME}^* &= \frac{k_{\text{off}}}{k_{\text{off}} + k_{\text{on}}} \\
\text{B}_0^* &= \frac{k_{\text{off}} k_{\text{on}}}{(k_{\text{off}} + k_{\text{on}})(k_{\text{off}} + k_{\text{t}})} \\
&\vdots \\
\text{B}_i^* &= \text{B}_0^* \alpha^i \\
&\vdots \\
\text{S}^* &= \frac{k_{\text{on}}}{k_{\text{off}} + k_{\text{on}}} \alpha^N
\end{aligned}$$

The steady state fraction of activated TCR complexes, $\text{S}^*$, can also be interpreted as the approximate fraction of time a single TCR, in a single simulation, spends in an activated state. The usefulness of this interpretation is that we can use $\text{S}^*$ to find the mean first passage time of TCR activation, given that a TCR and agonist are capable of binding. Let $t_c$ be the time it takes a single TCR/agonist pair to complete one circuit starting immediately after dissociation from an activated TCR state (ME state), and then making the passage back to an activated state (multiple dissociations are possible). Then

$$t_c = t_{\text{kp}} + t_{\text{S}}$$

The time, $t_{\text{kp}}$, is the mean time it takes for a TCR to negotiate the kinetic proofreading mechanism and become activated, given that an agonist is always in binding range. The mean includes the possibility of multiple dissociation events. The time $t_S = 1/k_{\text{off}}$ is the expected time an activated TCR remains in an activated state. The fraction of time a TCR is activated becomes

$$S^* = \frac{t_{\text{kp}}}{t_{\text{kp}} + t_S} \qquad [1]$$

Since $S^*$ is known and $t_S = 1/k_{\text{off}}$, we can solve for $t_{\text{kp}}$:

$$\begin{aligned} t_{\text{kp}} &= \frac{t_S(1 - S^*)}{S^*} \\ &= \frac{k_{\text{on}}(1 - \alpha^{\text{N}}) + k_{\text{off}}}{k_{\text{off}} k_{\text{on}} \alpha^{\text{N}}} \\ &= \frac{1}{k_{\text{off}}} \left( \frac{1}{\alpha^{\text{N}}} - 1 \right) + \frac{1}{\alpha^{\text{N}}} \frac{1}{k_{\text{on}}} \end{aligned}$$

The term, $1/k_{\text{off}}$, is the average dissociation time in the kinetic proofreading mechanism. The difference $(1/\alpha^{\text{N}} - 1)$ is the mean number of attempts necessary to successfully become activated minus the one successful attempt. Finally, $(1/\alpha^{\text{N}})(1/k_{\text{on}})$ is the expected number of TCR/Antigen binding events multiplied by the expected time for a TCR and agonist to bind.

### 2. Fitting a Generalised Logistic Function to T-cell Activation Data

We assume that T-cell activation in a population follows a binomial distribution with parameter $\theta$. If $X$ is the random variable associated with T-cell activation, then this gives the following model (given 100 independent trials),

$$X_i = \begin{cases} 0 & \text{T-cell does not activate} \\ 1 & \text{T-cell activates} \end{cases}$$

$$\sum_{i=1}^{100} X_i \sim \text{Bin}(100, \theta)$$

$$P(x) = \binom{100}{x} \theta^x (1 - \theta)^{100-x}$$

The simulation data is used to approximate $\theta$ for a model with a given set of fixed parameters. More specifically, we fit a Generalised Logistic Model (GLM) to T-cell activation data given an agonist population (in the interface) of $n$ and a varying agonist dissociation rate $k_{\text{off}}$, while holding all other parameters constant for each $(k_{\text{off}}, n)$. Hence, $\theta_n = \theta_n(k_{\text{off}})$, where $\theta_n$ is the binomial parameter corresponding to the probability of T-cell activation for a model with $n$ agonists in the interface. Our logistic model takes the form of

$$\theta_n(k_{\text{off}}) = A + \frac{K - A}{(C + \exp(-B * (k_{\text{off}} - M)))^{1/v}}$$

with parameters having the following properties:

- $A$ : Lower asymptote: $A = 0$ for all models.
- $B$ : A selectivity term. Large B indicates a sharp transition from low probability of activation to high probability of activation.
- $v$ : A term that becomes relevant for small populations of agonists.

- $A + \frac{K-A}{C^{1/v}}$ : The upper asymptote. At sufficiently large agonist populations, $C = v = K = 1$.

- $M$ : A parameter that indicates where the increase in probability is maximal, i.e., $\max(\frac{d\theta_n(k_{\text{off}})}{dk_{\text{off}}}) = \frac{d\theta_n(M)}{dk_{\text{off}}}$

Figure 2 shows the GLM fits for our MV scanning model with $k_s = 100s^{-1}$ and a simulated interface time of $t_{\text{sim}} = 300s$. The intersecting horizontal line shows the points corresponding to $\theta_{n_i}(k_{\text{off}}^{(i)}) \approx 0.15$, where $k_{\text{off}}^{(i)}$ is the dissociation rate of $n_i$ agonists in a T-cell/APC interface such that there is a 15% chance of model activation. The purpose of the non-linear fitting of the activation data is to find the points of intersection which correspond to the T-cell activation contour ($P_{15}$), and the linear slope, $\gamma$, can be used to compare discrimination powers among models.

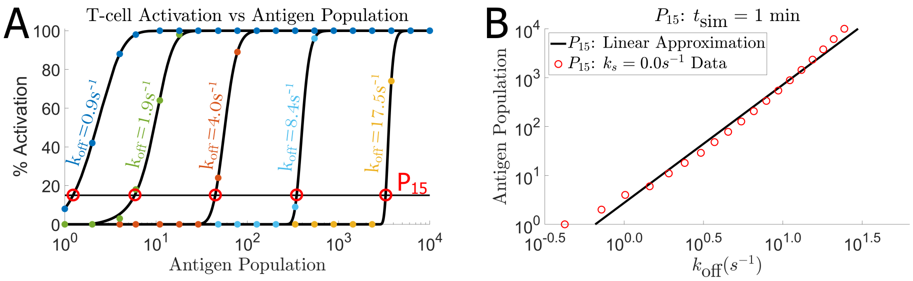


**Fig. 2.** Demonstration of the fitting method in order to compute $\gamma$. The example shown is a 3-step KP model with MV dynamics and $k_s = 100s^{-1}$. (**A**) Model Activation in a population of T-cells for a given agonist population on the APC contact surface and a varying dissociation rate, $k_{\text{off}}$. Points represent the actual activation data from simulations. The curves are fitted generalized logistic models (GLM). The black horizontal line represents $P_{15}$, and the intersection of this line with the fitted curves is the approximate point where each model achieves 15% T-cell activation in a population. The intersection points are then plotted and shown in the large agonist population region in (**B**). A linear model is then fitted to these points and the slope is the measure $\gamma$.

### 3. The Error in Approximating T-cell Activation Probability

We assume that for each dissociation rate and antigen population, $(k_{\text{off}}, n)$, that T-cell activation in a population follows a binomial distribution with parameter $\theta$. There are four conditions necessary for a random variable $X$ to follow a binomial distribution.

1) The first condition is that the experiment consists of $k$ identical trials. This condition holds in the probability sense for our simulations. While antigen and MV placement can vary from simulation to simulation, their probability distributions (for a given arrangement) are constant given a set of parameters.
2) Each outcome of T-cell activation in our model yields a binary result.
3) $\theta$ is constant for each set of parameters.
4) All $k$ trials are independent.

This yields the model:

$$X_i = \begin{cases} 0 & \text{T-cell does not activate} \\ 1 & \text{T-cell activates} \end{cases}$$

$$\sum_{i=1}^{k} X_i \sim \text{Bin}(k, \bar{\theta})$$

To illustrate this, we sample the number of activated T-cells in a sample population of 100, over M trials (the number of trials chosen is different among models). Our assumption is that the number of activated

T-cells in each sample population of 100 T-cells is distributed according to $\mathrm{Bin}(k, \bar{\theta})$. Therefore, we estimate $\bar{\theta}$ as

$$\bar{\theta} = \frac{1}{Mk} \sum_{j=1}^{M} \sum_{i=1}^{k} X_{ij} \qquad [2]$$

where $X_{ij}$ is the i-th sample of the j-th trial. The estimated distribution is shown along with sample data in Figure 3A–C for the three models with $k_{\mathrm{off}} = 1.0s^{-1}$ and $n = 1$ antigen.

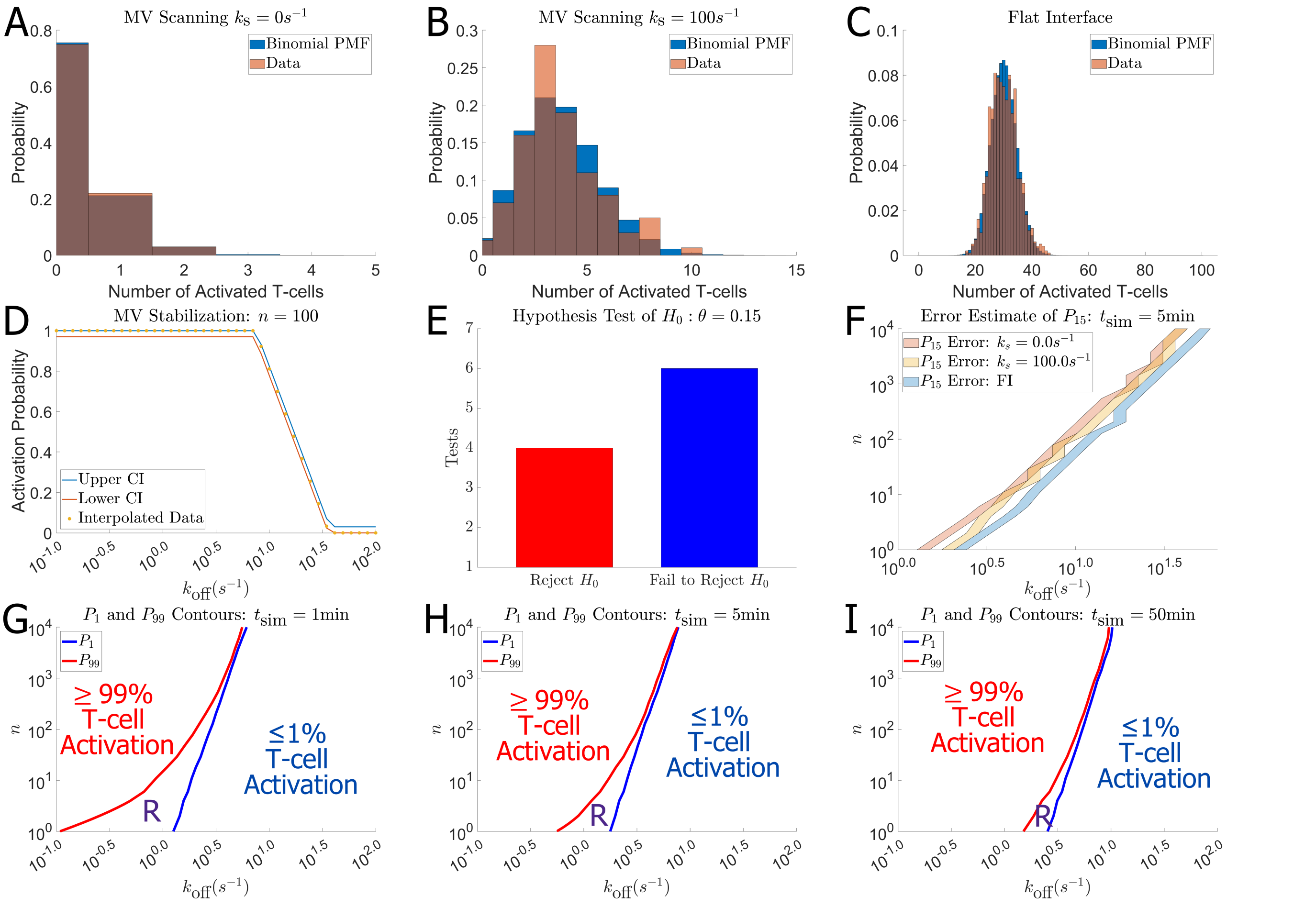


**Fig. 3.** The primary source of errors in the calculation of the discrimination power is the interpolated points from the T-cell activation data (e.g. $\mathrm{P}_{15}$). **A–C** For any fixed set of parameters, T-cell activation follows a binomial distribution with parameter $\theta$ which can be estimated using Eq. (2). A binomial PMF using this estimate, and data from simulations, is compared for the MV scanning (**A**), MV stabilization (**B**), and the FI (**C**) models. Each simulation consists of a single agonist in the interface with dissociation rate ($k_{\mathrm{off}} = 1.0s^{-1}$). **D** A confidence interval about T-cell activation data shown for the MV stabilization model with 100 agonists in the interface. The interval is derived using "Jeffrey's Prior" and the "Rule of Three". The interval shown was calculated for interpolated points as well. **E** Hypothesis test showing that the confidence intervals do not necessarily hold about interpolated points. Here, $H_0$ is the hypothesis that an interpolated point ($\theta \approx 0.15$) is within the confidence interval. **F** An error estimate about interpolated data corresponding to the potency contours of which the discrimination power is derived from. The error for each point is the distance between the two closest explicitly simulated points that encompass the interpolated point. This provides a greater than or equal to 95% confidence interval for interpolated points. **G–I** Plots showing that the area (region R) between potency contours decreases as T-cell/APC interface time increases. For many antigen populations, this means that the $k_{\mathrm{off}}$ interval over which all $0.01 < \theta < 0.99$ becomes more narrow. As time grows large, it is possible to have all such $\theta$ be within two explicitly simulated points, given an insufficiently small discretization of $k_{\mathrm{off}}$. This feature also has implications in terms of discrimination.

While $\theta$ may be constant at the macroscopic scale for a given set of conditions, it also contains the majority of the complexity involved in our model (i.e. MV scanning, antigen arrangement, kinetic proofreading mechanism etc.), and is not easily computed analytically. Therefore, we approximate $\theta$ by simulation and interpolate the data to get a continuous approximation. The approximation is of the form $\theta \approx \theta_n(k_{\mathrm{off}})$, where $n$ is the number of agonists in the interface and $k_{\mathrm{off}}$ is their dissociation rate from TCRs. We then utilize this approximation to locate the point at which we would expect to see a percentage, say 15%, of activated T-cells in a population, $\theta_{n_i}(k_{\text{off}}^{(i)}) \approx 0.15$, for a given antigen density $n_i$. The point, $(k_{\text{off}}^{(i)}, n_i)$, corresponding to $\theta_{n_i}(k_{\text{off}}^{(i)})$ is one of the data points used to fit the linear model described in "Fitting a Generalised Logistic Function to T-cell Activation Data". We then construct a 95% confidence interval on the explicitly simulated points used to construct the interpolation function, $\theta_n(k_{\text{off}})$ by utilizing Jeffrey's Interval and the "Rule of Three"(Figure 3D). For the data where $0 < \bar{\theta} < 1$, this entails using Jeffrey's Prior for the prior distribution of the mean, $\bar{\theta}_p \sim \text{Beta}(1/2, 1/2)$. The posterior is then computed as

$$\bar{\theta} \sim \text{Beta}(\sum_{i=1}^{k} X_i + 1/2, 100 - \sum_{i=1}^{k} X_i + 1/2) \qquad [3]$$

The 95% confidence interval is computed by finding the 0.025 and 0.975 quantiles of the posterior distribution. For $\bar{\theta} = 0$ or $\bar{\theta} = 1$, we use the "Rule of Three", which simply states that the approximate 95% confidence intervals are $(0, 3/k)$ for $\bar{\theta} = 0$ and $(1 - 3/k, 0)$ for $\bar{\theta} = 1$. We note that due to the properties of Binomial random variables, the error associated with the observed data is quite small. This implies that Eq. (2) is a reasonable approximation of the parameter $\theta$ in the cases where the conditions were explicitly simulated. However, this confidence interval does not apply to any interpolated points. It is these points that are the primary source of errors in our computation of the discrimination power.

The primary source of the errors in our measures is the approximation of the binomial parameter $\bar{\theta}$ found via interpolation of the explicitly simulated data (interpolation of the sample means found via Eq. (2)), that give a sample mean 15% chance of T-cell activation ($\bar{\theta} \approx 0.15\%$). We can qualitatively evaluate the errors at the interpolated points by utilizing a simple hypothesis test. In our framework, our null hypothesis is described as $H_0 : \{\bar{\theta} = 0.15\}$ and the alternative is such that $H_a : \{\bar{\theta} \neq 0.15\}$. We then explicitly simulate the interpolated points derived from our data (Figure 2A) and construct a 95% confidence interval about these points using the same method as described previously. If our interpolated $\bar{\theta}$ falls within this interval, we fail to reject our initial hypothesis that $\bar{\theta} \approx 0.15$. On the other hand, we reject the null hypothesis in favor of the alternative if $\bar{\theta}$ is not in the confidence interval. We visualize this method by counting the number of interpolated points where we reject $H_0$ and fail to reject $H_0$ for a given model (Figure 3E). In our discretization of the $(k_{\text{off}}, n)$ space, this yields 19 interpolated binomial parameters (19 simulated antigen populations) paired with 19 sample binomial parameters ($\bar{\theta}$). In many cases, we must reject our assumption that $\bar{\theta} \approx 0.15$. This indicates that our method of interpolation, and/or the resolution of our discretization, is insufficient to determine the points, $\text{P}_{15}$, via interpolation. However, we know that explicitly simulated points yield a reasonable approximation of $\bar{\theta}$ (Figure 3A–D). Therefore, a reasonable error interval about any interpolated points, such as those corresponding to $\text{P}_{15}$, would be the distance in $k_{\text{off}}$ from the closest surrounding explicitly simulated points. This simply gives an error interval which is the width of a single discretized cell on the $k_{\text{off}}$ axis. This error interval about $\text{P}_{15}$ is shown in Figure 3F for 5 minute long simulated interfaces. While the approximation of $\text{P}_{15}$ is subject to significant error, in general, these errors have little influence in the qualitative relationship among models. For example, in Figure 3F we see that the regions where $\text{P}_{15}$ likely resides overlaps between MV Models with and without contact stabilization when there is a large population of antigen on the APC. Conversely, these regions diverge given smaller populations of antigen. Additionally, the stabilized MV model yields regions for $\text{P}_{15}$ that are close in value to $\text{P}_{15}$ regions of the flat interface model. Thus, even given our consideration of the error in estimating $\text{P}_{15}$, we come to similar conclusions about the relationships among our stochastic models.

Lastly, we note an interesting property of kinetic proofreading models with a temporal activation (signal) component. Consider interpolating any point $\text{P}_x$ such that $0 < x < 100$ for a given antigen population $n$ (Inverse of Figure 2A). If there is some interval in $k_{\text{off}}$, $\text{I}_n$, in which contains all $\text{P}_x : x \in (0, 100)$ for a given antigen population $n$, then $\text{R}_n(t) = \text{I}_n^{(h)}(t) - \text{I}_n^{(l)}(t)$ is a monotonically decreasing function with respect to time, where $\text{I}_n^{(h)}(t)$ is the right boundary and $\text{I}_n^{(l)}(t)$ is the left boundary on the $k_{\text{off}}$ axis. When taken over a range of antigen values, this yields a 2-dimensional region R with an approximate discretized area of $\text{A}(t) = \sum_n (\text{I}_n^{(h)}(t) - \text{I}_n^{(l)}(t))$. This region is shown in Figure 3G–I. As the simulation time is increased, the area $\text{A}(t)$ between contours decreases. One implication of this is that if the discretization of the $k_{\text{off}}$ axis is insufficiently fine, then it is likely that given sufficiently long simulation time (or T-cell/APC interface time), all $P_x$ for any $0 < x < 100$ will appear nearly identical. Indeed, this is nearly the case in Figure 3I. Another interesting implication this feature has is the ties with antigen discrimination. In our work, we loosely quantify antigen discrimination by measuring the discrimination power of a single $P_{15}$ contour. However, discrimination would be more accurately quantified by the distance between a high probability contour, like $P_{99}$, and a low probability activation contour such as $P_1$. We might expect host antigen dissociation rates to fall to higher $k_{\text{off}}$ than $P_1$ (Right of blue line in Figure 3G–I), and agonist dissociation rates to fall to lower $k_{\text{off}}$ than $P_{99}$ (Left of red line in Figure 3G–I). In this case, discrimination would be classified by the area of the region R between the two contours, where large $A(t)$ would indicate poor discrimination. What we have shown in Figure 3D–F) is that increasing the simulation time (T-cell/APC contact time) results in a decrease in the area of region R, and hence improves a T-cell's discriminatory ability by this metric. This now gives two instances where increasing T-cell/APC contact time improves antigen discrimination, one by increasing the discrimination power of an individual contour, and the other by decreasing the distance between all contours.

### 4. Mass Action Contours and Convergence

The antigen potency, $\rho$, is the number of antigen needed in the interface to elicit an average number of activated TCRs at steady state ($\Psi$) for any TCR/antigen dissociation rate in an N-step kinetic proofreading model. This formulation yields $\Psi = nS^*$, where $n$ is the number of antigen in the interface and $S^* = S^*(k_{\text{off}})$ is the fractional steady state value corresponding to a population of activated TCRs (Supplement:Mass action kinetic proofreading model). Multiplying the fractional steady state solution by the number of antigen gives $nS^* = \Psi$ and

$$\Psi = n\left(\frac{k_{\text{on}}}{k_{\text{on}} + k_{\text{off}}}\right)\left(\frac{k_{\text{t}}}{k_{\text{off}} + k_{\text{t}}}\right)^N \quad [4]$$

where $k_{\text{on}}$, $k_{\text{t}}$, and $N$ are constant for a given model. If $\Psi$ is set as constant for all antigen populations and dissociation rates, then any increase in dissociation rate that decreases $S^*(k_{\text{off}})$ must also result in an increase in $n = n(k_{\text{off}})$ to maintain the constant activated TCR population $\Psi$, i.e., the antigen population and fractional steady state solution become functions of $k_{\text{off}}$. Therefore, defining $\rho$ as $\rho(k_{\text{off}}) = n(k_{\text{off}})$ gives

$$\rho(k_{\text{off}}) = \left(\frac{k_{\text{on}} + k_{\text{off}}}{k_{\text{on}}}\right)\left(\frac{k_{\text{off}} + k_{\text{t}}}{k_{\text{t}}}\right)^N \Psi \quad [5]$$

where $\rho$ is the antigen potency contour for a given constant $\Psi > 0$, and the subscript $c$ denotes that this is the steady state activated TCR population over the entire contour (for all $k_{\text{off}}$). In terms of Eq. (5), $\Psi$ can be interpreted as a term that defines the sensitivity condition of a model. An example is such that, if we assume a T-cell is sensitive to some steady state activated TCR population, $\Psi$, then $\rho(k_{\text{off}})$ describes the number of antigen needed in the interface to elicit a steady state population of $\Psi$ activated TCRs for any dissociation rate, i.e., achieve the required sensitivity of the T-cell ($\Psi = 3$ in Figure 4A).

The usefulness of defining $\rho$ in such a way becomes apparent when comparing T-cell activation potencies ($P_{15}$) to that of antigen potencies ($\rho$). In some cases, $P_{15}$ appears nearly identical to $\rho$, for some sensitivity condition $\Psi_{\text{M}}$ (Figure 4B and C). This applies not only to the flat interface model, which is most similar to that of the mass action kinetics, but also MV models. This indicates that there are regions of $(k_{\text{off}}, n)$ such that the stochastic implementation of our models, with or without MV scanning, yields nearly identical antigen potency contours to that of the mass action model that only considers the underlying kinetic proofreading mechanism. In other words, in these regions of T-cell activation $P_{15}(k_{\text{off}}) \approx \rho(k_{\text{off}})$ and

$$P_{15}(k_{\text{off}}) \approx \left(\frac{k_{\text{off}} + k_{\text{on}}}{k_{\text{on}}}\right)\left(\frac{k_{\text{t}} + k_{\text{off}}}{k_{\text{t}}}\right)^N \Psi_{\text{M}} \quad [6]$$

Where $\Psi = \Psi_{\text{M}}$ and $M = \{\text{FI}, \text{MV}, \text{Stab}\}$. The term $\Psi_{\text{M}}$ has an additional meaning to $\Psi$ in Eq. (5), which is that $\Psi_{\text{M}}$ is the T-cell sensitivity condition required to give a 15% chance of T-cell activation. The potency,

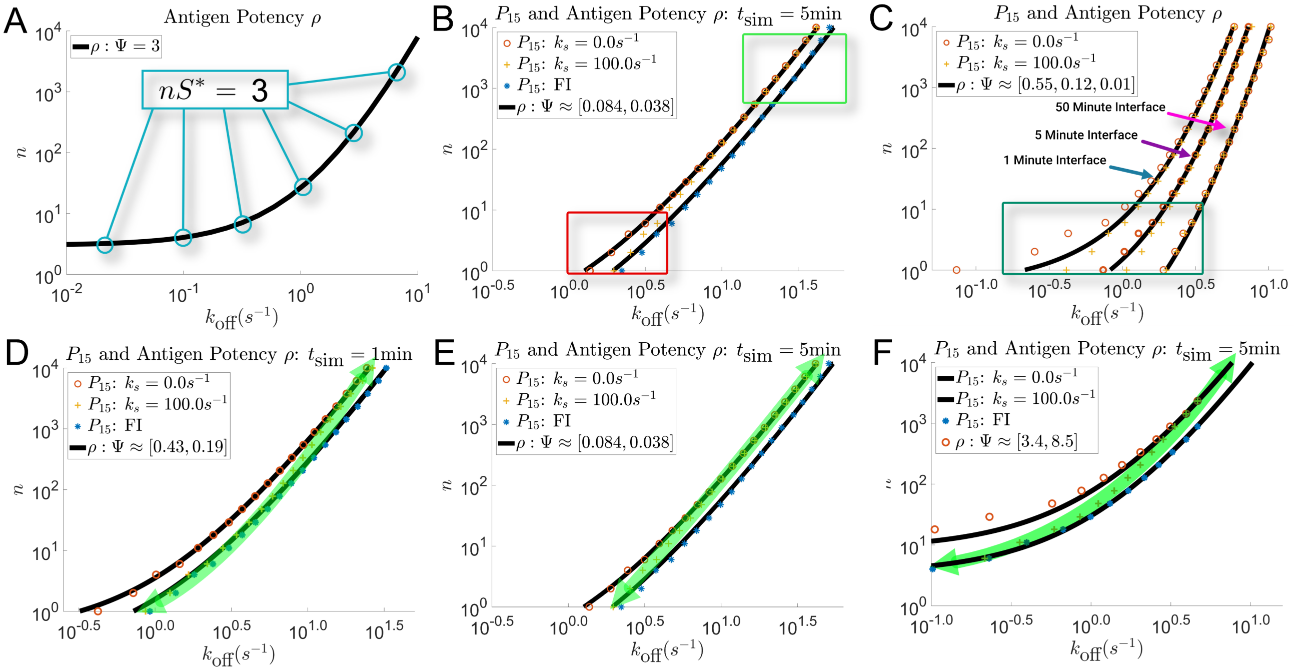


**Fig. 4.** Comparisons of T-cell activation contours with that of antigen potency contours. **A** Antigen potency contour where $\Psi = 3$ in Eq. (5). For any $k_{\text{off}}$, $\rho$ gives the number of antigen needed in the interface to elicit 3 activated TCRs at steady state. **B** T-cell activation contours ($P_{15}$) from data corresponding to a MV model without stabilization, MV stabilization model, and a flat interface model, with a simulated T-cell/APC interface time. Each model has identical kinetic proofreading parameters. Black curves are antigen potencies ($\rho$) with the same kinetic proofreading parameters, but different constant valued ($\Psi$) contours, where the specific values are denoted in the legend. The green box denotes a region where Eq. (6) is true. Conversely, the red boxed data shows a region where Eq. (6) is not true. **C** Data from MV models, with and without MV stabilization, in simulations of a T-cell and APC in contact for 1 minute, 5 minutes, and 50 minutes. Increasing the interface time increases the range of antigen populations where Eq. (6) is true. Furthermore, this shows an example of how $\Psi = \Psi(t)$ is decreases as interface time increases in Eq. (6). **D–F** Example data sets that show the property of MV stabilization that often gives higher discrimination powers than the flat interface model. The green arrows highlight $P_{15}$ from MV stabilization models joining $P_{15}$ from MV models without stabilization in large $n$ to $P_{15}$ of flat interface models at small $n$.

$P_{15}$, has an identical meaning to $\rho$, which is that $P_{15}$ is the number of antigen necessary to elicit the steady state activated TCR population of $\Psi_{\text{M}}$ for any $k_{\text{off}}$. This approximate convergence only applies under certain conditions, which are subject to change given different model parameters. In general, we observe that Eq. (6) is a valid approximation for sufficiently large antigen populations in the T-cell/APC interface (Figure 4B). However, what constitutes "sufficiently large" is model dependent and can be biologically infeasible. Additionally, we observe Eq. (6) in T-cell activation data following sufficiently long-lasting T-cell/APC contacts (Figure 4C). Similarly, what constitutes sufficiently long is model dependent. Together, our results indicate that increasing the number of antigen in the T-cell/APC interface, or increasing the time that a T-cell and APC are in contact, will increase the "likeness" of $P_{15}$ to $\rho$, where $\rho$ is defined by some T-cell sensitivity condition $\Psi_{\text{M}}$.

In regions of T-cell activation where Eq. (6) is observed for multiple stochastic models, such as the green boxed region in Figure 4B, $P_{15}$ derived from a MV scanning model only differs from the flat interface model by the sensitivity condition $\Psi_{\text{M}}$ (given identical KP parameters), with $\Psi_{\text{FI}} < \Psi_{\text{MV}}$. As described above, if $\Psi_{\text{M}}$ is treated as a parameter describing a model's sensitivity, then this would indicate that a flat interface model is more sensitive to antigen in the T-cell/APC contact. This is true in our stochastic models simply because of the additional geometric and TCR activation constraints imposed by MV scanning. In addition, we observe that MV scanning models with no contact stabilization mechanism ($k_{\text{s}} = 0s^{-1}$) and MV models with a high rate of contact stabilization ($k_{\text{s}} = 100s^{-1}$) become indistinguishable in regions where Eq. (6) is true (Figure 4B and C). Hence, when Eq. (6) is observed in a region, MV models with or without MV stabilization yield the same discrimination powers and sensitivity conditions ($\Psi_{\text{MV}} \approx \Psi_{\text{Stab}}$), in said region. In sufficiently long-lasting T-cell/APC contacts, this region can encompass all discrete antigen populations

(Figure 4C: 50 minute interface).

For simplicity, we classify T-cell activation data being representative of early or late activation phases by using Eq. (6). If $\Psi_{\mathrm{M}}$ is constant for all $k_{\mathrm{off}}$ with $n \geq 1$, then this represents late-phase T-cell activation. Conversely, if $\Psi_{\mathrm{M}}$ is constant only for sufficiently large antigen populations, then this is characterized as an early T-cell activation phase. In terms of Figure 4C, only the data from a 50 minute T-cell/APC contact would be characterized as a late phase activation event, while the 1 and 5 minute T-cell/APC interface data would be characterized as early phases. Using this characterization, our results indicate that in some early phases of T-cell activation, MV models with the additional feature of contact stabilization yield higher discrimination powers than flat interface models and MV models without the chance of contact stabilization. The higher discrimination power is the result of T-cell activation potencies derived from MV stabilization models merging with those of MV models without stabilization in large antigen populations ($\Psi_{\mathrm{Stab}} = \Psi_{\mathrm{MV}}$ for large $n$) and merging with those of flat interface models in smaller antigen populations ($\Psi_{\mathrm{Stab}} = \Psi_{\mathrm{FI}}$ for small $n$). This feature allows MV stabilization models to have a similar antigen sensitivity to flat interface models when there are only a few agonists in the interface (green arrows in Figure 4A–C). Since flat interface models will always be more sensitive to antigen than MV models (No MV search or removal), then this is an increase in antigen sensitivity as a result of the stabilization mechanism. The discrimination power is also increased because of MV models with stabilization having nearly identical T-cell activation contours as MV models without stabilization, when antigen populations are sufficiently large. However, the increased discrimination power is only temporary, and only present in certain comparable intervals. As the time that the T-cell and APC are in contact, or the number of antigen in the interface increases, the MV stabilization model becomes more similar to that of a MV model without stabilization (Figure 4C). As noted previously, in late stages of T-cell activation MV models with and without stabilization become indistinguishable ($\Psi_{\mathrm{Stab}} \approx \Psi_{\mathrm{MV}}$ for all $n$ in Eq. (6)). If MV and flat interface models are both in the late stage of T-cell activation, then in any comparable interval ($n \geq 1$) the flat interface model will have the higher discrimination power. This is a result of Eq. (6), and is described in the section titled "Increasing T-cell Sensitivity to Antigen also Increases the Discrimination Power".

Finally, we show that any model parameter that improves a T-cell's sensitivity to antigen without changing the underlying properties of the kinetic proofreading mechanism, will also increase the discrimination power of said model in all comparable regions where $\Psi_{\mathrm{M}}$ is constant. An example of such a comparable region wold be the antigen populations encompassed by the green box in Figure 4B, and the comparisons that could be made would be the discrimination power over these antigen populations for each model. In said region, the flat interface would yield the higher discrimination power due to being more sensitive to antigen ($\Psi_{\mathrm{FI}} < \Psi_{\mathrm{MV}}$). Another such example of a parameter that could accomplish this would be $k_{\mathrm{act}}$, which amplifies the signal from activated TCRs. In our stochastic model, $k_{\mathrm{act}}$ amplifies the signal from activated TCRs that can lead to T-cell activation. A larger $k_{\mathrm{act}}$ (Figure 4E) would give smaller values of $\Psi_{\mathrm{M}}$ versus that of a smaller $k_{\mathrm{act}}$ (Figure 4F). Thus, in antigen populations where $\Psi_{\mathrm{M}}$ is constant, increasing $k_{\mathrm{act}}$ increases the discrimination power. The other parameter in our stochastic models that has this affect is the T-cell/APC contact time, $t_{\mathrm{sim}}$. It is obvious that increasing the time a T-cell and APC are in contact will increase the sensitivity to antigen if there is no signal decay. However, it is less obvious that this will also increase the discrimination power, with this increase being significant under certain conditions. Hence, increasing the sensitivity of T-cells to antigen in the T-cell/APC interface, without changing any properties of the underlying kinetic proofreading mechanism of TCR/antigen interactions, also increases the discrimination power of the T-cell given the condition that $\Psi_{\mathrm{M}}$ is constant in the measured regions. This is a counter intuitive result as it does not reflect the well known property of kinetic proofreading models, which is that increases in the discrimination power (increasing $N$) are accompanied with a worsened sensitivity to antigen. However, it can be proven using the antigen potency contours.

### 5. Increasing T-cell Sensitivity to Antigen also Increases the Discrimination Power

If $\Psi_{\mathrm{M}}$ is constant for some stochastic model, M, in some region, then a T-cell activation contour (such as $P_{15}$) in said region is nearly identical to some antigen potency contour $\rho$ with the sensitivity condition $S_{\mathrm{c}}^{*} = \Psi_{\mathrm{M}}$. Let Model A and Model B be two T-cell activation models with identical kinetic proofreading mechanisms in which this condition is met in some comparable region, such as the flat interface model and MV models in the green boxed region of Figure 4B. Let $\rho_{\mathrm{A}}$ and $\rho_{\mathrm{B}}$ be the antigen potency contours for which the T-cell activation contours from Model A and Model B have approximately converged (such as the solid black curves in Figure 4B), respectively. Furthermore, assume that $S_{\mathrm{c}}^{*} = \Psi_{\mathrm{A}}$ and $S_{\mathrm{c}}^{*} = \Psi_{\mathrm{B}}$ are the respective sensitivity conditions, where $\Psi_{\mathrm{A}} > \Psi_{\mathrm{B}}$. In our interpretation of $\Psi_{\mathrm{M}}$, this indicates that Model B is more sensitive to antigen in the T-cell/APC interface. Our goal is to show that any equivalent change in dissociation rate ($\epsilon$), given an equivalent initial potency (Eq. (7)), will yield a larger change in antigen potency in Model B (i.e., a higher discrimination power). An equivalent initial potency forces the condition that

$$\rho_{\mathrm{B}}(k_{\mathrm{off}}^{(\mathrm{B})}) = \rho_{\mathrm{A}}(k_{\mathrm{off}}^{(\mathrm{A})}) \qquad [7]$$

where $k_{\mathrm{off}}^{(\mathrm{B})} > k_{\mathrm{off}}^{(\mathrm{A})}$ because $\Psi_{\mathrm{A}} > \Psi_{\mathrm{B}}$ and all kinetic proofreading parameters being equivalent. Thus, to accomplish our goal it suffices to show

$$\rho_{\mathrm{B}}(k_{\mathrm{off}}^{(\mathrm{B})} + \epsilon) - \rho_{\mathrm{B}}(k_{\mathrm{off}}^{(\mathrm{B})}) \geq \rho_{\mathrm{A}}(k_{\mathrm{off}}^{(\mathrm{A})} + \epsilon) - \rho_{\mathrm{A}}(k_{\mathrm{off}}^{(\mathrm{A})}) \qquad [8]$$

where

$$\rho_{\mathrm{B}} = \left(\frac{k_{\mathrm{off}} + k_{\mathrm{on}}}{k_{\mathrm{on}}}\right)\left(\frac{k_{\mathrm{t}}}{k_{\mathrm{t}} + k_{\mathrm{off}}}\right)^{-N} \Psi_{\mathrm{B}}$$

$$\rho_{\mathrm{A}} = \left(\frac{k_{\mathrm{off}} + k_{\mathrm{on}}}{k_{\mathrm{on}}}\right)\left(\frac{k_{\mathrm{t}}}{k_{\mathrm{t}} + k_{\mathrm{off}}}\right)^{-N} \Psi_{\mathrm{A}}$$

Taking the derivative of $\log_{10}[\rho]$ with respect to $\log_{10}[k_{\mathrm{off}}]$ gives

$$\frac{d\log_{10}[\rho]}{d\log_{10}[k_{\mathrm{off}}]} = k_{\mathrm{off}}\left(\frac{1}{k_{\mathrm{on}} + k_{\mathrm{off}}} + \frac{N}{k_{\mathrm{t}} + k_{\mathrm{off}}}\right) \qquad [9]$$

which is a monotonically increasing function with respect to $k_{\mathrm{off}}$ (or $\log_{10}[k_{\mathrm{off}}]$). Therefore,

$$\log_{10}[\rho_{\mathrm{B}}(k_{\mathrm{off}}^{(\mathrm{B})} + \epsilon)] - \log_{10}[\rho_{\mathrm{B}}(k_{\mathrm{off}}^{(\mathrm{B})})] \geq \log_{10}[\rho_{\mathrm{A}}(k_{\mathrm{off}}^{(\mathrm{A})} + \epsilon)] - \log_{10}[\rho_{\mathrm{A}}(k_{\mathrm{off}}^{(\mathrm{A})})] \qquad [10]$$

since $k_{\mathrm{off}}^{(\mathrm{B})} > k_{\mathrm{off}}^{(\mathrm{A})}$ and the derivative at every point, $\log_{10}[\rho_{\mathrm{B}}(k_{\mathrm{off}}^{(\mathrm{B})} + \epsilon^{(\mathrm{i})})]$ will be larger than the corresponding point, $\log_{10}[\rho_{\mathrm{A}}(k_{\mathrm{off}}^{(\mathrm{A})} + \epsilon^{(\mathrm{i})})]$, for $\epsilon^{(\mathrm{i})} \in [0, \epsilon]$. From Eq. (10), we can derive

$$\frac{\rho_{\mathrm{B}}(k_{\mathrm{off}}^{(\mathrm{B})} + \epsilon)}{\rho_{\mathrm{B}}(k_{\mathrm{off}}^{(\mathrm{B})})} \geq \frac{\rho_{\mathrm{A}}(k_{\mathrm{off}}^{(\mathrm{A})} + \epsilon)}{\rho_{\mathrm{A}}(k_{\mathrm{off}}^{(\mathrm{A})})} \qquad [11]$$

Using Eq. (7) gives $\rho_{\mathrm{B}}(k_{\mathrm{off}}^{(\mathrm{B})} + \epsilon) \geq \rho_{\mathrm{A}}(k_{\mathrm{off}}^{(\mathrm{A})} + \epsilon)$. We can then subtract both sides by $\rho_{\mathrm{B}}(k_{\mathrm{off}}^{(\mathrm{B})})$ and $\rho_{\mathrm{A}}(k_{\mathrm{off}}^{(\mathrm{A})})$, respectively, which gives the inequality Eq. (8).

In regions where $\Psi_{\mathrm{M}}$ is constant, from Eq. (8) we know that any decrease in $\Psi_{\mathrm{M}}$ will increase the discrimination power of a given model, up to $\gamma = N + 1$. Additionally, we know there are many parameters that can modulate $\Psi_{\mathrm{M}}$ without changing the kinetic proofreading mechanism, such as MV dynamics or the signal amplification parameter, $k_{\mathrm{act}}$. Perhaps the most interesting result of Eq. (8) is we now see how

T-cell/APC interface time can modulate antigen discrimination in T-cells. All other parameters being fixed for a given model, just by increasing the time that a T-cell and APC are in contact can improve antigen discrimination and sensitivity. Intuitively, the increase in T-cell/APC contact time allows for weaker agonists to generate similar T-cell activation signals as stronger agonists in shorter T-cell/APC contact times (dependent on signal decay). Additionally, this leads to a distinction of $\Psi_{\mathrm{M}}$ from $S_{\mathrm{c}}^{*}$ in Eq. (5) and Eq. (6), where $S_{\mathrm{c}}^{*}$ is a constant in Eq. (5) and $\Psi_{\mathrm{M}} = \Psi_{\mathrm{M}}(t)$ is a decreasing function with respect to time in Eq. (6).

We can extrapolate from Eq. (8) a more mathematical description of the results in early activation phases shown in Figure 4. Firstly, there are early activation phases where all stochastic models out perform the expected discrimination power of regional antigen potency contours (data not shown). While $\Psi_{\mathrm{M}}$ will still be constant for sufficiently large antigen populations, this indicates that in the regions where $\Psi_{\mathrm{M}}$ is not constant, the discrimination power and antigen sensitivity will be increased due to these stochastic effects. In terms of modeling the $P_{15}$ contours, we should introduce a decaying function with respect to antigen population and time such that Eq. (6) becomes

$$P_{15}(k_{\mathrm{off}}, n, t) \approx \left(\frac{k_{\mathrm{off}} + k_{\mathrm{on}}}{k_{\mathrm{on}}}\right)\left(\frac{k_{\mathrm{t}} + k_{\mathrm{off}}}{k_{\mathrm{t}}}\right)^{N}(\Psi(t) - \delta_{\mathrm{stoch}}(n,t)) \qquad [12]$$

with $\delta_{\mathrm{stoch}}(n,t) > 0$ and $\delta_{\mathrm{stoch}}(n,t) \to 0$ as $n$ and/or $t$ increases. Since $(\Psi(t) - \delta_{\mathrm{stoch}}(n,t)) < \Psi(t)$, from Eq. (8) we know that this will increase the discrimination power of stochastic models in early T-cell activation phases, which is what we observe in the data. For a flat interface model, this would be the only modification necessary to model the T-cell activation contours with respect to time, in the mean sense. Thus, $\Psi_{\mathrm{FI}} = \Psi(t) - \delta_{\mathrm{stoch}}(n,t)$

To extend Eq. (12) to a MV scanning model we add another decaying term with respect to time and antigen population, which gives

$$P_{15}(k_{\mathrm{off}}, n, t) \approx \left(\frac{k_{\mathrm{off}} + k_{\mathrm{on}}}{k_{\mathrm{on}}}\right)\left(\frac{k_{\mathrm{t}} + k_{\mathrm{off}}}{k_{\mathrm{t}}}\right)^{N}(\sigma\Psi(t) - \delta_{\mathrm{stoch}}(n,t) + \delta_{\mathrm{MV}}(n,t)) \qquad [13]$$

where $\delta_{\mathrm{MV}}(n,t) > 0$ and $\delta_{\mathrm{MV}}(n,t) \to 0$ as $n$ and $t$ increase. The constant $\sigma$ is a scaling factor that accounts for the reduced sensitivity (increase in $\Psi$) caused by geometric and TCR activation constraints due to MV scanning. This means that even as the additional terms in Eq. (13) decay, the MV model will always have a discrimination power less than the flat interface model. This is an intuitive result, as the MV scanning model will always have an additional constraint to TCR/antigen interactions (MV contact removal $k_{\mathrm{d}}$), which does not exist in a flat interface model. Therefore, the MV model will always be less sensitive to antigen in the interface. Hence,

$$\lim_{n\to\infty}[\Psi_{\mathrm{MV}}(t)] = \sigma\Psi(t) \geq \Psi(t) = \lim_{n\to\infty}[\Psi_{\mathrm{FI}}(t)]$$
$$\lim_{t\to\infty}[\Psi_{\mathrm{MV}}(t)] = \sigma\Psi_{\infty} \geq \Psi_{\infty} = \lim_{t\to\infty}[\Psi_{\mathrm{FI}}(t)]$$

where $\Psi_{\mathrm{MV}} = \sigma\Psi(t) - \delta_{\mathrm{stoch}}(n,t) + \delta_{\mathrm{MV}}(n,t)$ and $\Psi_{\infty}$ is a possible limit imposed by kinetic proofreading parameters or activation signal decay. It is important to note that the rapidity of which the additional terms decay are dependent on model parameters. In the case of Figure 4C, we see that $\delta_{\mathrm{stoch}}(n,t) \to 0$ faster than $\delta_{\mathrm{MV}}(n,t) \to 0$. Conversely, in Figure 4D–E we observe $\delta_{\mathrm{stoch}}(n,t) \to 0$ slower than $\delta_{\mathrm{MV}}(n,t) \to 0$.

Finally, $P_{15}$ for MV stabilization models must also be modified with the addition of one more term. For stabilized MV models,

$$P_{15}(k_{\mathrm{off}}, n, t) \approx \left(\frac{k_{\mathrm{off}} + k_{\mathrm{on}}}{k_{\mathrm{on}}}\right)\left(\frac{k_{\mathrm{t}} + k_{\mathrm{off}}}{k_{\mathrm{t}}}\right)^{N}(\sigma\Psi(t) - \delta_{\mathrm{stoch}}(n,t) + \delta_{\mathrm{MV}}(n,t) - \delta_{\mathrm{stab}}(n,t)) \qquad [14]$$

where $\delta_{\text{stab}}(n,t) > 0$ and $\delta_{\text{stab}}(n,t) \to 0$ as $n$ or $t$ increases. Another property of $\delta_{\text{stab}}(n,t)$ is that $\delta_{\text{stab}}(n,t) \to 0$ as $t$ decreases. This is also an intuitive property. If the T-cell/APC interface time is shorter than $1/k_{\text{d}}$, then MV removal will become less probable and MV stabilization becomes redundant. Therefore, since $\delta_{\text{stab}}(n,t) > 0$, there likely exists a time $t$ where $\delta_{\text{stab}}(n,t)$ takes a maximal value. Lastly, we note that

$$\lim_{n\to\infty}[\Psi_{\text{stab}}(t)] = \sigma\Psi(t) = \lim_{n\to\infty}[\Psi_{\text{MV}}(t)]$$
$$\lim_{t\to\infty}[\Psi_{\text{stab}}(t)] = \sigma\Psi_{\infty} = \lim_{t\to\infty}[\Psi_{\text{MV}}(t)]$$

where $\Psi_{\text{stab}} = \sigma\Psi(t) - \delta_{\text{stoch}}(n,t) + \delta_{\text{MV}}(n,t) - \delta_{\text{stab}}(n,t)$. This formulations agrees with our stochastic data and observations that MV models with or without contact stabilization become indistinguishable in large antigen populations or later phases of T-cell activation.

### 6. MV Scanning and Packing in the T-cell/APC Interface

In our MV scanning model, MV contacts are equivalent in size and cannot overlap other MV contacts or the T-cell/APC boundary. These circular contacts are placed within the T-cell/APC interface, which is modeled as a disk with radius $5\mu$m. The placement of MV contacts is implemented by choosing two randomly drawn real numbers, $u, v \sim U[0,1]$, such that $\theta = 2\pi u$ and $r_{\text{center}} = R_{\text{IS}}\sqrt{v}$, where $R_{\text{IS}} = 5\mu$m and the MV contact is placed with its center at coordinates $(r_{\text{center}}, \theta)$. In order to implement the stochastic Gillespie Algorithm, we construct a propensity function for the placement of MV contacts in the T-cell/APC interface, which requires knowledge of the total number of MV contacts (applicable MV population) that can be placed. To approximate this number, we find an approximate packing limit of MV contacts in the T-cell/APC interface that can be achieved by our random contact placement method. This is accomplished by setting a threshold number of trials, $T$, such that if a packing simulation fails to place an additional contact $T$ times, then the simulation has reached a random packing limit and the final number of MV contacts are recorded ($A_{\text{M}}$). The final number of contacts found using this method can vary considerably with respect to MV contact sizes (Figure 6A). However, the final area coverage is often similar, given these large variations. The final area coverage is computed as $n_{\text{MV}}r_{\text{MV}}^2/R_{\text{IS}}^2$, where $n_{\text{MV}}$ is the number of MV contacts and $r_{\text{MV}}$ is the MV contact radius (Figure 6B). It is important to note that this method fails to approximate an optimal packing limit, as we can see in Figure 6C that the likelihood of finding an available space to place an additional MV contact can be dependent on $T$. Additionally, we see this is increasingly true with smaller MV contact radii. The reason we utilize this method given the apparent shortcomings is that the packing limit, and the method by which the packing limit is achieved, is largely unknown in vivo.

The instantaneous area coverage in MV scanning simulations (Figure 6D–F) was recorded at each time point identically to the method above ($n_{\text{MV}}r_{\text{MV}}^2/R_{\text{IS}}^2$). We set the mean instantaneous area coverage to approximately 40%. To accomplish this, we determine a $k_{\text{a}}$ such that,

$$k_{\text{a}} = \frac{k_{\text{d}}n_{\text{MV}}^*}{A_{\text{M}} - n_{\text{MV}}^*} \qquad [15]$$

where $A_{\text{M}}$ is the maximum number of contacts that can be placed in the interface for a given MV radius ($r_{\text{MV}}$), and $n_{\text{MV}}^*$ is the number of MV contacts required to cover 40% of the interface. In Figure 6D, varying the MV contact radius also changes the maximum number of contacts that can be placed in the interface and the number of contacts required to cover 40% of the T-cell/APC interface. Thus, $k_{\text{a}}, n_{\text{MV}}^*$, and $A_{\text{M}}$ also vary with MV radius. In Figure 6E, varying $k_{\text{d}}$ only leads to a change in $k_{\text{a}}$ to maintain the 40% area coverage. In Figure 6F, none of the parameters in Eq. (15) are changed. However, the instantaneous area fraction increases as a result of MV contacts stabilizing. As MV contacts stabilize, fewer contacts are subject to removal, which leads to a larger number of contacts in the interface. This results in an increase in the instantaneous area fraction, with the extreme being that all contacts are stabilized, and the number of contacts in the interface is $A_{\text{M}}$.

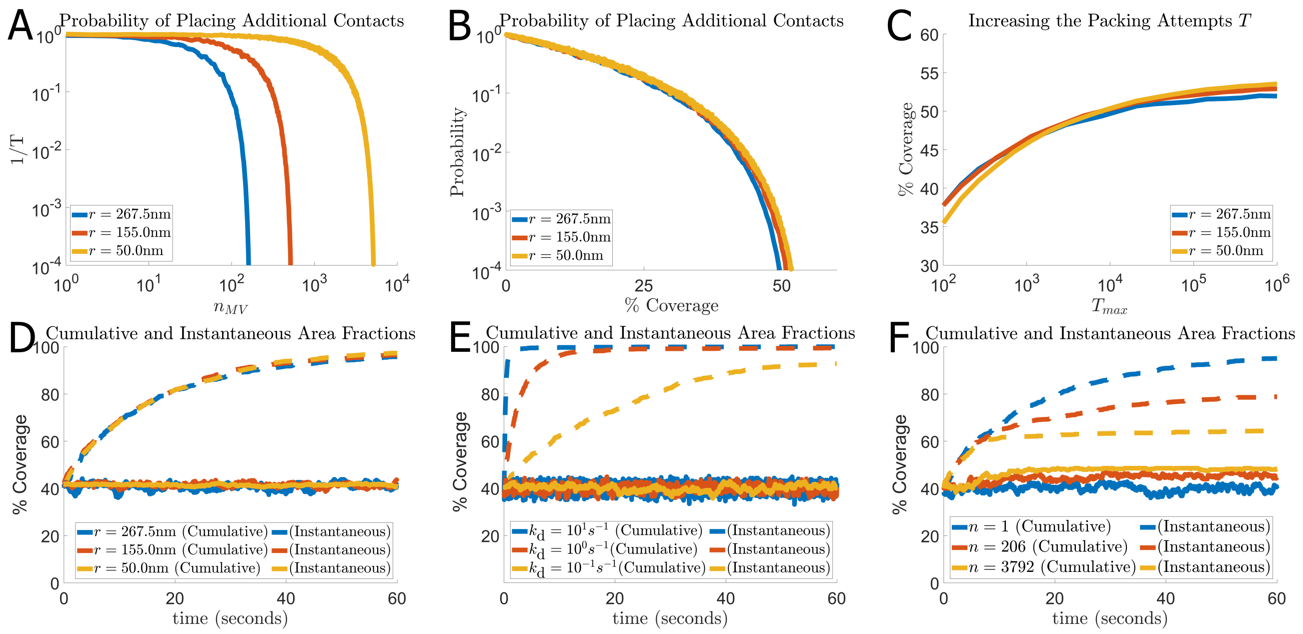


**Fig. 5**

**Fig. 6.** Packing limits obtained by random contact placement and cumulative/instantaneous scanning fractions of MV. **A** The probability of placing an additional contact when there are $n_{\mathrm{MV}}$ contacts in the interface. Decreases in the MV contact radius can have a significant effect on the maximum number of contacts placed in the interface (horizontal intercept). **B** Probability of placing an additional contact with respect to the interface area covered by existing contacts. While the number of contacts can vary significantly, the final area coverage by contacts is often similar. **C** Final area coverage of contacts with respect to the number of attempts alloted to successfully place an additional contact. We note that increasing the number of attempts, $T_{\max}$, also increases the final area coverage. While increasing $T_{\max}$ will eventually lead to a horizontal asymptote, there can still be notable increases in the final area coverage even for large $T_{\max}$. **D** Cumulative and instantaneous scanning fractions over time for a varied MV radius, $r_{\mathrm{MV}}$. The cumulative scanning speed is only slightly increased as MV radius is decreased. Additionally, smaller MV radii exhibit smaller fluctuations about the mean instantaneous area coverage. **E** Cumulative and instantaneous scanning coverage for a varied $k_{\mathrm{d}}$. If the instantaneous area coverage is maintained constant (Eq. (15)), then increasing $k_{\mathrm{d}}$ also increases the cumulative scanning speed of MV contacts. **F** Cumulative and instantaneous area fractions in the presence of agonists. Agonists can cause MV contacts to become stabilized, which in turn causes a slower cumulative scanning and a larger instantaneous area fraction. In the extreme case, where nearly all MV contacts have become stabilized, the instantaneous area coverage approaches a packing limit.

The cumulative area fraction is the total T-cell/APC interface area visited by at least one contact at time $t$. This is found by using an algorithmic method known as Vogel's Method, which is implemented by generating coordinates $(\tau_i, p_i)$ in a disk such that

$$p_i = i\theta$$

$$\tau_i = \sqrt{\frac{i}{M}}$$

for all $i \leq M$, where $\theta$ is the golden angle and $M$ is the number of points approximately equally spaced in the disk. The cumulative area fraction $(CAF(t))$ can then be approximated by counting the number of points that have been covered by at least one MV contact at time t, $m(t)$, divided by the total number of points. Hence,

$$CAF(t) = m(t)/M \qquad [16]$$

Using the described MV dynamics, we observe that changing the MV contact radius can increase scanning speed, but only slightly (Figure 6D). A parameter with a more significant influence on scanning speed is $k_{\mathrm{d}}$. Increasing $k_{\mathrm{d}}$ increases the scanning speed of MV contacts, given Eq. (15). As previously explained, the instantaneous area fraction is maintained at approximately 40%. When $k_{\mathrm{d}}$ is increased, MV contacts are being removed at a faster rate. Likewise, to maintain the constant mean instantaneous area fraction, $k_{\mathrm{a}}$ must also increase. Hence, increasing $k_{\mathrm{d}}$ increases the flux of MV contact removal, which increases the flux of MV contact placement, given that the mean instantaneous area fraction is constant. In a random MV contact placement method, this results in an increase in the scanning speed, $dm(t)/dt$, i.e., an increase in the cumulative scanning fraction at time $t$ (Figure 6E). Conversely, MV stabilization slows the cumulative scanning speed by decreasing the flux of both, MV contact placement contact removal. As more contacts are stabilized by antigen interaction, the number of contacts in the interface at any point in time increases. This decreases the propensity function for which contacts are placed in the interface, $k_{\mathrm{a}}(A_{\mathrm{M}} - n_{\mathrm{MV}})$. Stabilized MV contacts are not subject to removal from the interface, which also decreases the propensity function for MV removal ($k_{\mathrm{d}} n_{\mathrm{MV}}$). Hence, we see that the strength and number of foreign antigen can influence MV scanning speeds by stabilizing contacts (Figure 6F).

### 7. Dynamics of MV Lifetimes

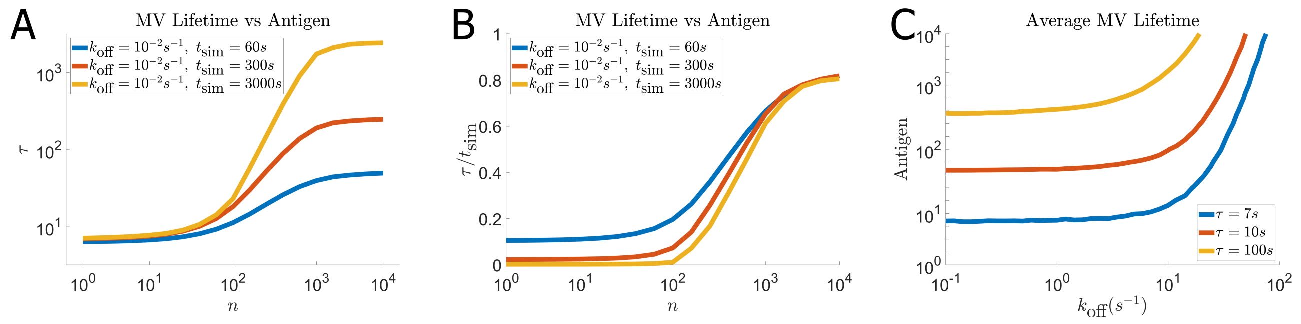


**Fig. 7.** Average MV contact lifetimes given varying antigen populations and agonist potency. **A** Average lifetimes for MV in the presence of a varied population of strong agonists ($k_{\mathrm{off}} = 10^{-2}s^{-1}$) and three different T-cell/APC interface times. When there is a strong agonist presence, the time over which MV contact times are recorded can significantly impact the average. This is due to many MV contacts remaining until the simulation is terminated (or observation is complete). **B** By dividing the average MV contact times by the total observation time ($\tau/t_{\mathrm{sim}}$), we observe an approximate convergence for multiple simulation times when there are large populations of strong agonists in the interface. This is a key indicator that the obtained average MV contact lifetimes are heavily weighted by existing contacts that remain until the simulation is terminated. **C** Average MV contact times with respect to a varying $k_{\mathrm{off}}$ and antigen population, $n$. MV contact lifetimes obtained from our simulations are highly dynamic in nature. If MV are rarely stabilized, the average approaches $1/k_{\mathrm{d}}$ (below blue contour). This is true in small populations of antigen, even given a strong agonist. As agonist populations increase, so too does the average MV contact lifetimes (given sufficient agonists strength).

In a MV model without stabilization, we set the average MV contact lifetime to $1/k_{\mathrm{d}} = 7s$. The average MV lifetime in a simulation of the T-cell/APC interface is

$$\tau = \sum_{i=1}^{M} \tau_i + \sum_{j=1}^{Q} \tau_j \qquad [17]$$

where $\tau_k$ is the dwell time of the $k^{th}$ contact, $M$ denotes the number of contacts that have been removed, and $Q$ denotes the number of contacts that still exist when the simulation is terminated. When antigen can stabilize MV contacts, the average MV lifetime can be heavily weighted by the summation $\sum_{j=1}^{Q} \tau_j$. In general, this applies to cases where the average MV lifetime is close to, or greater than, the T-cell/APC interface time ($\tau \geq t_{\mathrm{sim}}$). In Figure 7A, this would be the case for $n \geq 10^4$, since these are simulations with strong agonists. Clearly, this is not an approximation of the average MV lifetime in these extremes. A way we can determine this is the case from data is by recording the average MV lifetime given extreme conditions (small $k_{\mathrm{off}}$ and large $n$) and then dividing by the total time over which the T-cell/APC contact was observed. In the case of our stochastic model, this would be $\tau/t_{\mathrm{sim}}$. In the case where MV have longer average contact times than the simulation period, we observe that this fraction converges to a constant for multiple interface times, $t_{\mathrm{sim}}$ ($n \approx 10^4$ in Figure 7B). However, in smaller antigen populations, or weaker agonists, This fraction is not constant and decreases as the T-cell/APC interface time increases. These observations indicate that recorded average MV lifetimes can be influenced not only by parameters, antigen potency, and antigen quantity, but also the total time over which the data as observed.

1. TW McKeithan, Kinetic proofreading in t-cell receptor signal transduction. *Proc. Natl. Acad. Sci.* **92**, 5042 (1995).
